# RIP140 inhibits glycolysis-dependent proliferation of breast cancer cells by regulating GLUT3 expression through transcriptional crosstalk between hypoxia induced factor and p53

**DOI:** 10.1101/2020.07.30.228759

**Authors:** Valentin Jacquier, Delphine Gitenay, Samuel Fritsch, Sandrine Bonnet, Balázs Győrffy, Stéphan Jalaguier, Laetitia K. Linares, Vincent Cavaillès, Catherine Teyssier

**Author notes:** Corresponding authors (C.T.); (V.C.). IRMB, University of Montpellier, INSERM, CNRS, CHU Montpellier, Montpellier, France. Senior authors (Vincent Cavaillès, Catherine Teyssier).

## Abstract

Cancer cells with uncontrolled proliferation preferentially depend on glycolysis to grow, even in the presence of oxygen. The transcriptional co-regulator RIP140 represses the activity of transcription factors that drive cell proliferation and metabolism and plays a role in mammary tumorigenesis. Here we use cell proliferation and metabolic assays to demonstrate that RIP140-deficiency causes a glycolysis-dependent increase in breast tumor growth. We further demonstrate that RIP140 reduces the transcription of the glucose transporter GLUT3 gene, by inhibiting the transcriptional activity of hypoxia inducible factor HIF-2α in cooperation with p53. Interestingly, RIP140 expression was significantly associated with good prognosis only for breast cancer patients with tumors expressing low GLUT3, low HIF-2α and high p53, thus confirming the mechanism of RIP140 anti-tumor activity provided by our experimental data. Overall, our work establishes RIP140 as a critical modulator of the p53/HIF cross-talk to inhibit breast cancer cell glycolysis and proliferation.

## Introduction

In a normal resting cell, glycolysis converts glucose to pyruvate, which enters the tricarboxylic acid cycle where it becomes oxidized to generate ATP into mitochondria. In the absence of oxygen, glucose is still degraded into pyruvate, which is now converted into lactate. On switching to proliferative mode, cells increase glycolysis and reduce oxidative phosphorylation, which results in a high rate of glycolysis leading to lactate production. This metabolic switch was first described by Otto Warburg in the 20’s, who observed that cancer cells prefer glycolysis to mitochondrial respiration to produce energy, even in the presence of oxygen [1]. This conversion of glucose to lactate is now established as the Warburg effect, or aerobic glycolysis. At first, Warburg hypothesized that cancer arises from impaired mitochondria. However, experimental observations of functional mitochondria in cancer cells have since refuted this hypothesis [2]. In fact, increased aerobic glycolysis has been frequently verified in tumors. It is now established that cancer cells switch to glycolysis to supply their increased need for biosynthetic precursors and organic resources to synthetize cell components [3]. As glycolysis produces less ATP than oxidative phosphorylation, cancer cells compensate for this energy gap by up-regulating glucose transporters to import more glucose into the cell. Enhanced glycolysis also produces reducing equivalents and many glycolytic intermediates are diverted into the pentose phosphate pathway to produce NADPH [4], allowing cancer cells to fight against reactive oxygen species and oxidative stress [5].

The switch to glycolysis in cancer cells is orchestrated by oncogenes and tumor suppressors [6]. Specifically, the tumor suppressor gene p53 favors oxidative phosphorylation over glycolysis, enhancing the production of reactive oxygen species, to imbalance redox status and promote cell death [7, 8]. p53 directly inhibits glycolysis by blocking the expression of glucose transporters [9, 10]. Conversely, inactivating p53 reduces oxygen dependence and allows cancer cells to grow in oxygen-limited conditions, such as hypoxia. Hypoxia induces the stability of hypoxia inducible factors, or HIFs, which have an oxygen sensitive α sub-unit (HIF-1α, HIF-2α or HIF-3α) and one stable β sub-unit (HIF-1β; also known as ARNT) [11]. In cancer cells, HIFs antagonize p53 and stimulate glycolysis. However, the mechanisms of their direct influence on glycolysis needs further clarification [12, 13].

The transcriptional coregulator RIP140, also called NRIP1 (Nuclear Receptor Interacting Protein 1) regulates the activity of various transcription factors, mainly inhibiting their transactivation ability by recruiting histone deacetylases[14]. RIP140 can also activate transcription through SP1 [15], or NF-κB [16] sites. Due to its wide interactome, RIP140 influences numerous physiological functions, such as mammary gland development, fertility and inflammation [17]. RIP140 is also a major regulator of energy homeostasis, regulating lipogenesis in the liver, lipid storage and thermogenesis in adipose tissues, mitochondrial integrity, and the formation of oxidative fibers in the muscle [17]. In normal adipocytes, RIP140 has been shown to reduce the expression of GLUT4 [18] and its trafficking at the cell membrane [19, 20].

RIP140 plays crucial roles in solid tumors [21]. In colorectal cancers, RIP140 regulates the APC/β-catenin pathway and inhibits the proliferation of intestinal cells. Moreover, RIP140 acts as a tumor suppressor, high expression of RIP140 correlating with good prognosis of colorectal cancer patients [22, 23]. In breast cancer, we, and other groups, have reported that RIP140 promotes [24–26] or impairs mammary tumor cell proliferation [27, 28]. Whether RIP140 is involved in the glucose metabolism of cancer cells and, if so, whether this directly affects tumor proliferation is still unknown.

Here we simultaneously characterized the effects of RIP140 on proliferation and glycolysis of cancer cells. We knocked-down RIP140 expression in human breast cancer cells and used immortalized and transformed mouse embryonic fibroblasts from RIP140 knockout (RIPKO) mice. We evaluated the survival of RIP140-deficient cells after disrupting glycolysis and characterized the mechanisms by which RIP140 regulates glycolysis in breast cancer cells to influence their growth. In line with these data, RIP140 expression correlates with good prognosis in breast cancer patients defined by a signature of the newly identified target gene and transcription partners.

## Results

### RIP140-deficiency promotes cell proliferation and tumor growth

As mentioned above, opposite effects of RIP140 on human breast cancer cell proliferation have been described. Aiming to clarify this situation, we evaluated the impact of RIP140 silencing after small interfering RNA (siRNA) knockdown with two separate siRNAs on the proliferation of MCF7 and MDA-MB-436 breast cancer cell lines.

As shown by xCELLigence real-time cell analysis and MTT assays (Fig. 1a, b and Supplementary Fig. 1a-e), RIP140 silencing consistently increased cell proliferation confirming our previous results [27, 28]. To further validate the antiproliferative activity of RIP140 on human cancer cells, we performed the same type of experiments in prostate (DU145) and colon (RKO) human cancer cell lines. As shown in Supplementary Fig. 1e, knocking-down RIP140 expression in these cancer cells robustly increased proliferation in our experimental conditions.

**Fig. 1.**
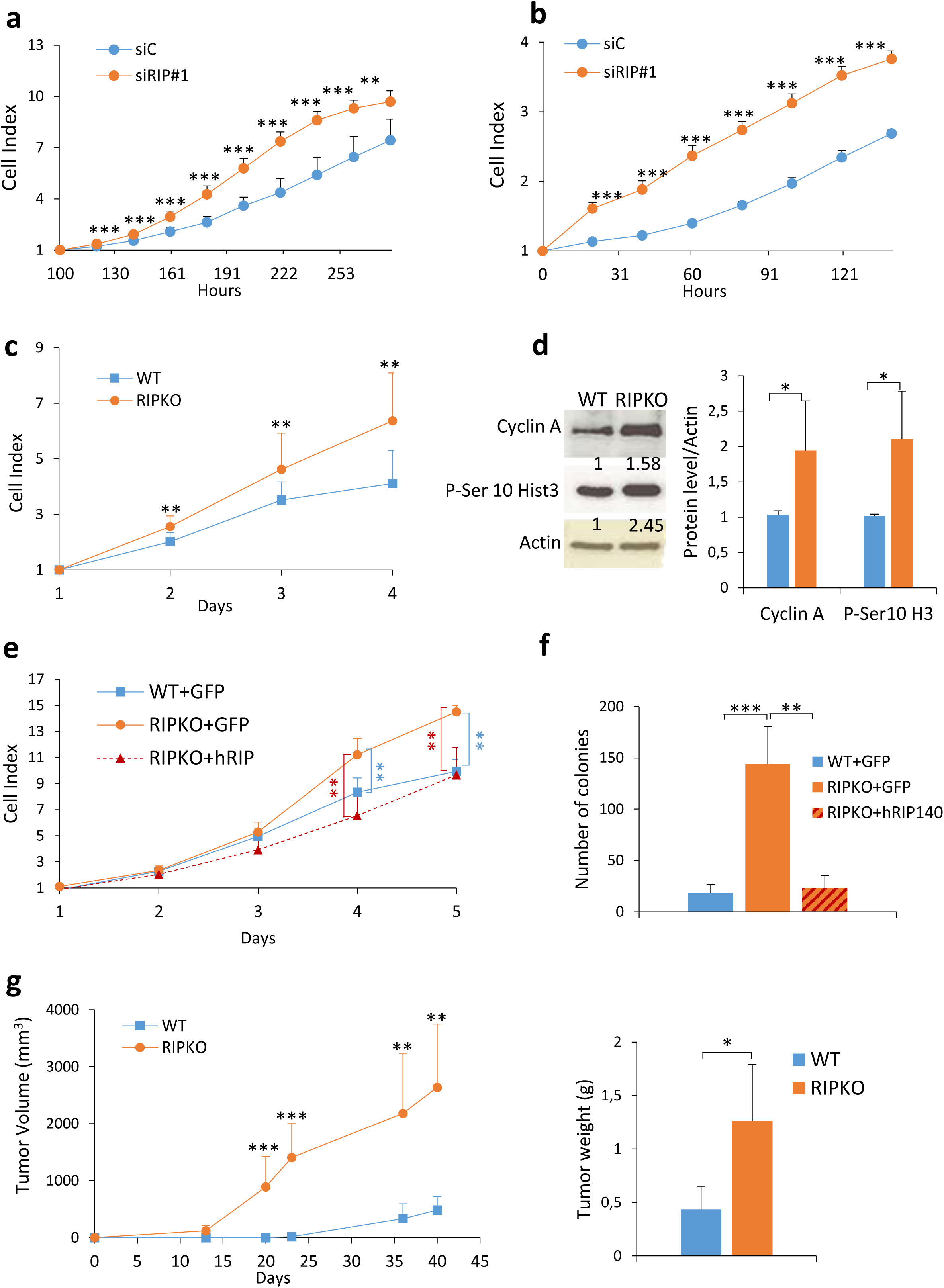
RIP140-deficiciency enhances cell proliferation and tumorigenesis. **a, b**, MCF7 (**a**) and MDA-MB-436 (**b**) cells were transfected with control siRNA (siC) or RIP140 siRNA (siRIP#1). Live measurements of cell proliferation were performed with the xCELLigence RTCA DP instrument. The cell index values (average and standard deviation) of three independent experiments are shown. For the MCF7 cells, the cell index was normalized to 100h to account for slow proliferation rate. (**p < 0.01, ***p < 0.001). **c**, Live measurements of cell proliferation were performed with the xCELLigence RTCA DP instrument in immortalized MEFs (MEF #1) WT and RIP140 KO (RIPKO) cells (mean ± SD, n=3, ** p <0.01). **d**, *Left panel*: Representative blot of Cyclin A and Phospho-Serine 10 Histone H3 protein concentration assessed by Western blot analysis in MEF #1 cells. *Right panel*: Densitometry analysis by normalizing to the housekeeper Actin and represented as a mean of n = 3 ± SD (*p <0.05). Unprocessed original scans of blots are shown in Supplementary Figure 7. **e**, Live measurements of cell proliferation were performed with the xCELLigence RTCA DP instrument in MEF #1 cells stably overexpressing a pEGFP plasmid WT (WT+GFP) or RIPKO (RIPKO+GFP) and RIPKO stably overexpressing a pEGFP-hRIP140 plasmid (RIPKO+hRIP140). The statistical analysis depicted in blue characters compared the differences in the values between RIPKO+GFP and WT+GFP. The statistical analysis depicted in red characters compared the differences in the values between RIPKO+GFP and RIPKO+hRIP140 (mean ± SD, n=3, ** p <0.01). **f**, Number of colonies of H-RasV12-transformed MEF #1 cells stably overexpressing pEGFP or pEGFP-hRIP140 obtained in a colony formation in soft agar assay (mean ± SD, n=4, ** p <0.01, ***p <0.001). **g**, Nude mice xenograft experiments. SV40/H-RasV12-transformed MEF (MEF #2) WT and RIPKO cells were injected subcutaneously into the flanks of immuno-deficient mice (n=7) to assess their tumorigenic potential (left panel). The weight of the tumors was assessed six weeks after grafting (n=6 mice; right panel). (*p <0.05, **p <0.01, ***p <0.001).

We also studied the proliferation of mouse embryonic fibroblasts (MEFs) knocked-out for the *Rip140* gene (RIPKO, Supplementary Fig. 1f). 3T3-immortalized RIPKO MEFs proliferated more than 3T3-immortalized MEFs expressing *Rip140* (WT), as shown by xCELLigence and MTT assays (Fig. 1c and Supplementary Fig. 1g). Immortalized RIPKO cells expressed more the proliferation markers, Cyclin A and Phosphorylated Serine10-Histone 3, and were richer in ATP (Fig. 1d and Supplementary Fig. 1h). In SV40-HRas/V12-transformed MEFs, RIP140-deficiency increased the cell proliferation and the number of colonies as shown, respectively, by MTT and soft agar assays (Supplementary Fig. 1i, j). To confirm the role of RIP140 in the observed effects, we rescued RIP140 expression by generating stable MEFs expressing either GFP or human RIP140 (Supplementary Fig. 1k). Rescuing RIP140 expression inhibited cell proliferation (Fig. 1e and Supplementary Fig. 1l), and reduced colony number (Fig. 1f and Supplementary Fig. 1m).

As the above data built a strong case for a growth inhibitory role of RIP140, we wondered whether RIP140 could also affect tumor growth. We therefore xenografted transformed RIPKO MEFs into nude mice and found that, indeed, these tumors had enhanced growth when compared to transformed MEFs expressing *Rip140* (WT) (Fig. 1g and Supplementary Fig. 1n, o). Altogether, our findings strongly push toward an anti-proliferative effect of RIP140 on cancer cell proliferation.

### Inhibiting glycolysis reduces the growth advantage of RIP140-deficient cells

As glucose homeostasis is a key parameter in the control of tumor cell proliferation, we sought to determine the importance of glucose for RIP140-deficient cell growth. We first performed glucose starvation experiments and monitored cell proliferation. Lowering glucose concentrations affected RIPKO cell proliferation more than WT cells (Fig. 2a and Supplementary Fig. 2a). As shown in Supplementary Fig. 2b, the growth advantage of RIPKO cells observed in glucose-rich medium was abolished in the absence of glucose.

**Fig. 2.**
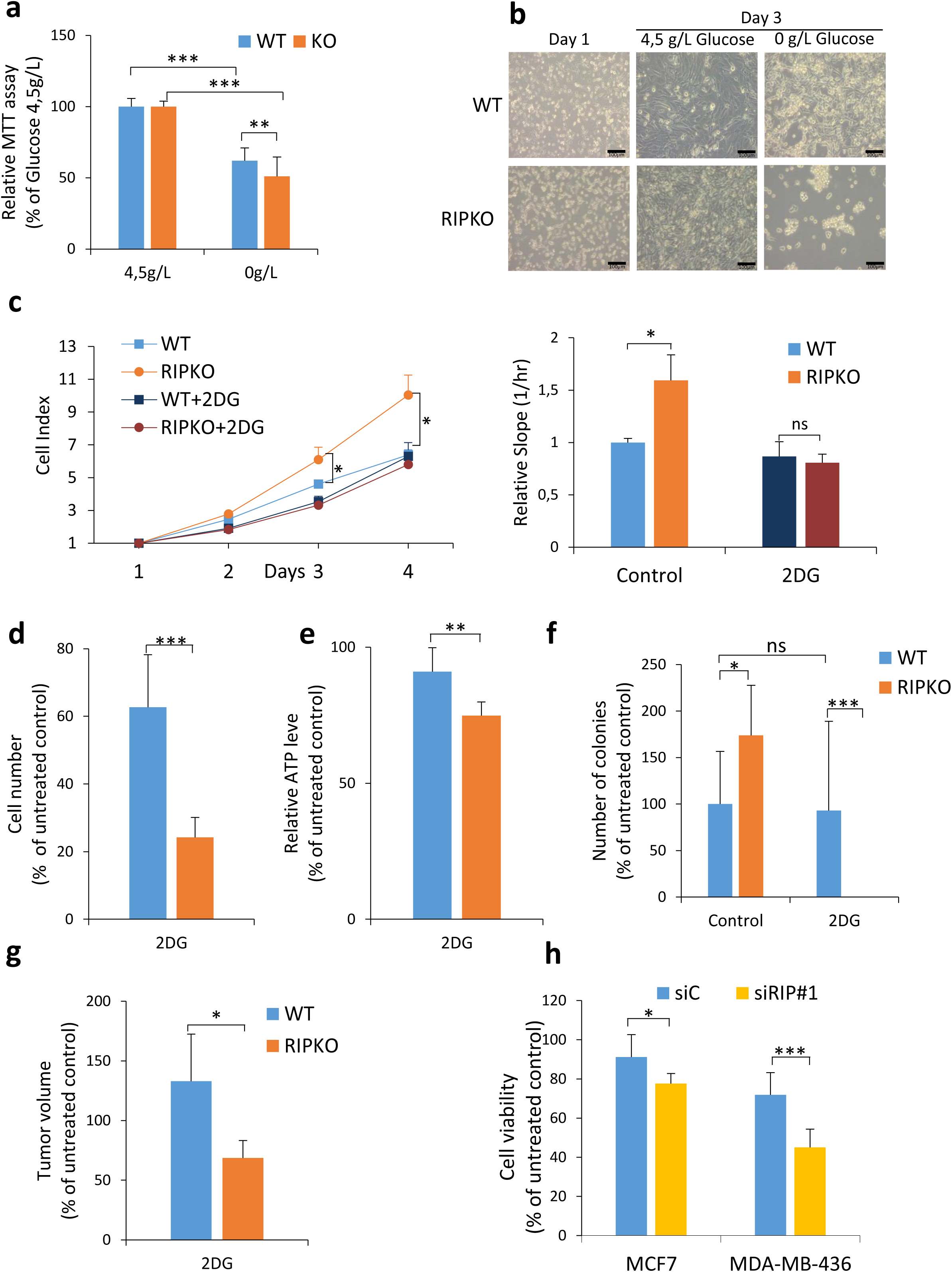
Inhibiting glycolysis reduces the growth advantage of RIP140-deficient cells. **a**, 3-(4,5-dimethylthiazol-2-yl)-2,5-diphenyltetrazolium Bromide (MTT) assay in MEF #1 cells cultured in DMEM-containing 0g/L or 4,5g/L glucose for 10 days. Results were normalized to the cell density in DMEM-containing 4.5g/L glucose for each cell line (mean ± SD, n=3, **p <0.01, ***p <0.001). **b**, Microscopic morphological analysis of SV40/H-RasV12-transformed MEF #3 cells cultured in DMEM-containing 0g/L or 4,5g/L glucose. Scale bar: 100µm. **c**, Live measurements of cell proliferation were performed with the xCELLigence RTCA DP instrument in MEF #1 cells treated or not with 2-deoxyglucose (2DG 5mM). The statistical analysis compared the differences in the values between untreated RIPKO and WT samples (left panel). The slopes were extracted using the xCELLigence RTCA Software from the curves in the left panel. Values are normalized to untreated WT cells (right panel). (mean ± SD, n=3, *p <0.05, *ns* not significant) **d**, Cell proliferation was assessed by counting MEF #1 cells treated or not with 2DG (5mM). Data are normalized to untreated control for each cell line after eight days of 2DG treatment (mean ± SD, n=3, ***p <0.001). **e**, Cellular ATP content in MEF #1 cells after 3 days of 2DG treatment (5mM). Values are normalized to that of untreated control for each cell line (mean ± SD, n=3, ***p <0.001). **f**, Colony formation in soft agar assay of transformed MEF #3 cells after 3 weeks of 2DG treatment (5mM). Number of colonies are normalized to untreated WT samples (mean ± SD, n=3, *p <0.05, ***p <0.001, *ns* not significant) **g**, Tumor volume of transformed MEF #1 cells xenografted in nude mice after eighteen days of 2DG administrated intra-peritoneally every other day (20mg/g). Data are normalized to that of untreated control for each cell line (mean ± SD, n=6, *p <0.05). **h**, Cell viability assessed by crystal violet staining of MCF7 and MDA-MB-436 cells transfected with control siRNA (siC) or RIP140 siRNA (siRIP#1) and treated for seven days with 2DG (5mM). Values are normalized to that of untreated control siRNA (mean ± SD, n=3, *p <0.05, ***p <0.001).

Upon glucose starvation, previous work reported that transformed MEFs displayed morphological features of cell death, such as loss of plasma membrane integrity and cell fragmentation (Chiaradonna et al., 2006). Transformed RIPKO MEFs displayed more features of dead cells than transformed WT cells (Fig. 2b). Furthermore, we found that transformed RIPKO MEFs were more sensitive to glucose limitation than transformed WT MEFs (Supplementary Fig. 2c). These data suggested that RIP140-deficiency triggered glucose starvation-induced cell death.

Next, we analyzed cell viability after glycolysis blockade with the hexokinase inhibitor 2-deoxyglucose (2DG). As for glucose starvation, 2DG treatment abolished the RIPKO cell growth advantage (Fig. 2c and Supplementary 2d). Moreover, treatment with the GAPDH inhibitor 3-Bromopyruvate (BrP) induced the same response (Supplementary Fig. 2e). 2DG treatment reduced cell proliferation and ATP level more in RIPKO MEFs (Fig 2d, e and Supplementary Fig. 2f). Moreover, RIPKO cells formed no colonies in the presence of the drug (Fig. 2f and Supplementary Fig. 2g). In preclinical models, to treat engrafted nude mice with 2DG led to a greater reduction in volume of RIPKO than WT tumors (Fig 2g). Finally, both glycolysis inhibitors reduced the viability of MCF7 and MDA-MB-436 cells more efficiently when RIP140 was silenced by siRNA (Fig. 2h and Supplementary Fig. 2h).

Altogether, these results demonstrate that the growth advantage of RIP140-deficient cells is abolished when glycolysis is impaired. Therefore, RIP140-deficient cells rely on glycolysis to grow, suggesting that RIP140 inhibits cell proliferation by blocking glycolysis.

### RIP140-deficiency enhances aerobic glycolysis

Because these results suggested that RIP140 inhibits glycolysis in tumor cells, we characterized the glycolytic properties of RIP140-deficient cells. We first demonstrated that glucose uptake was higher in immortalized and transformed RIPKO MEFs (Fig. 3a). We then performed Seahorse flux analysis that allow the measurement of the glucose-induced extracellular acidification rate, reflecting glycolysis. The glycolytic parameters, such as glycolysis, glycolytic capacity and glycolysis reserve, were higher in RIPKO MEFs (Fig. 3b and Supplementary Fig. 3a). The high glycolysis content of RIPKO cells was confirmed in transformed MEFs (Supplementary Fig. 3b). Furthermore, glucose consumption and lactate production were higher in RIPKO cells, confirming the Seahorse analysis (Fig. 3c, d). Then, rescuing RIP140 expression reduced extracellular acidification, confirming the inhibitory effect of RIP140 on glycolysis (Fig. 3e). Finally, silencing RIP140 by siRNA in MCF7 and MDA-MB-436 human breast cancer cells increased glycolysis (Fig. 3f and Supplementary 3c). Similar results were obtained in prostate and colon cancer cell lines (Supplementary Fig. 3d).

**Fig. 3.**
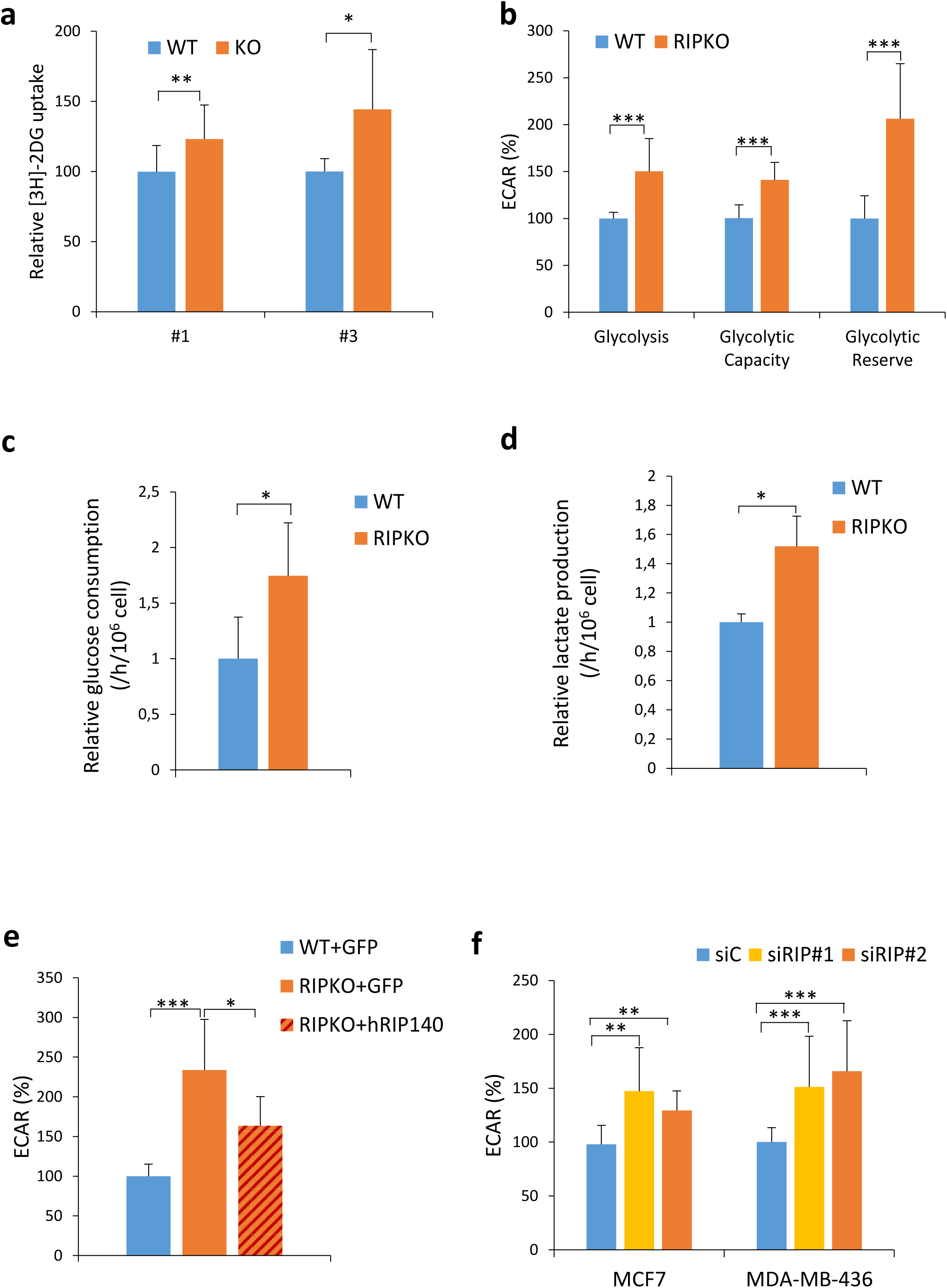
RIP140-deficiency enhances aerobic glycolysis. Unless otherwise stated MEF #1 were used. **a**, Glucose uptake assessed by [^3^H] 2-Deoxy-D-glucose uptake in immortalized MEF #1 (n=4) and transformed MEF #3 cells (n=3). Values are normalized to protein quantity and to the WT samples (mean ± SD, *p < 0.05, **p < 0.01). **b**, ECAR (Extracellular acidification rate) was measured in MEF #1 cells using the Seahorse XF96 analyzer. Different parameters of glycolytic functions (glycolysis=glucose-basal, glycolysis capacity=oligomycine-glucose, glycolytic reserve=glycolysis-glycolysis capacity), were calculated employing the Seahorse XF Glycolysis Stress Test. Values are normalized to basal measurements and to WT samples (mean ± SD, n=3, ***p < 0.001). **c**, **d**, Glucose consumption (**c**) and lactate production (**d**) were measured in MEF #1 cells by enzymatic determination assay. Cells were seeded in culture dishes and cultured for 8 h. The culture medium was collected and then changed. Cells were incubated for an additional 16h and counted. Medium samples were collected and used in enzymatic determination assay. Values were normalized to cell number, time unit and to WT samples (mean ± SD, n=3, *p < 0.05). **e**, Extracellular acidification rate (ECAR) measured using the Seahorse XF96 analyzer in MEF # 1 stably overexpressing a pEGFP plasmid WT (WT+GFP), RIPKO (RIPKO+GFP) and RIPKO stably overexpressing a pEGFP-hRIP140 plasmid (RIPKO+hRIP140). Values are normalized to basal measurements and to WT samples (mean ± SD, n=3, *p < 0.05, ***p < 0.001). **f**, Extracellular acidification rate (ECAR) measured using the Seahorse XF96 analyzer in MCF7 and MDA-MB-436 cells transfected with control siRNA (siC) or RIP140 siRNA (siRIP#1). Values are normalized to basal measurements and to control siRNA samples. The statistical analysis compared the differences in the values between control siRNA (siC) and each RIP140 siRNA (mean ± SD, n=3, **p< 0.01, ***p < 0.001).

Thus, taken together, these results demonstrate that the loss of RIP140 increased glycolysis in cancer cells.

### GLUT3 is essential for the growth advantage of RIP140-deficient cells

To investigate how RIP140 regulates glycolysis in cancer cells, we monitored by RT-qPCR the expression of 23 different metabolic genes involved in glycolysis, in WT and RIPKO immortalized MEFs. The mRNA levels of *Glut3*, *Glut4* and *Ldhb* (Lactate Dehydrogenase B) were increased more than 2-fold in RIPKO MEFs (Supplementary Fig. 4a). The increase of *Glut3* mRNA expression level was the most important and was recapitulated in transformed MEFs (Supplementary Fig. 4b). Furthermore, this gene was also significantly increased by RIP140 silencing in the breast cancer cell lines MCF7 and MDA-MB-436 (Fig. 4a and Supplementary Fig. 4c), whereas the *LDHB* and *GLUT4* levels did not change significantly (data not shown). Furthermore, rescuing RIP140 expression in RIPKO MEFs down-regulated *Glut*3 expression (Fig. 4b).

**Fig. 4.**
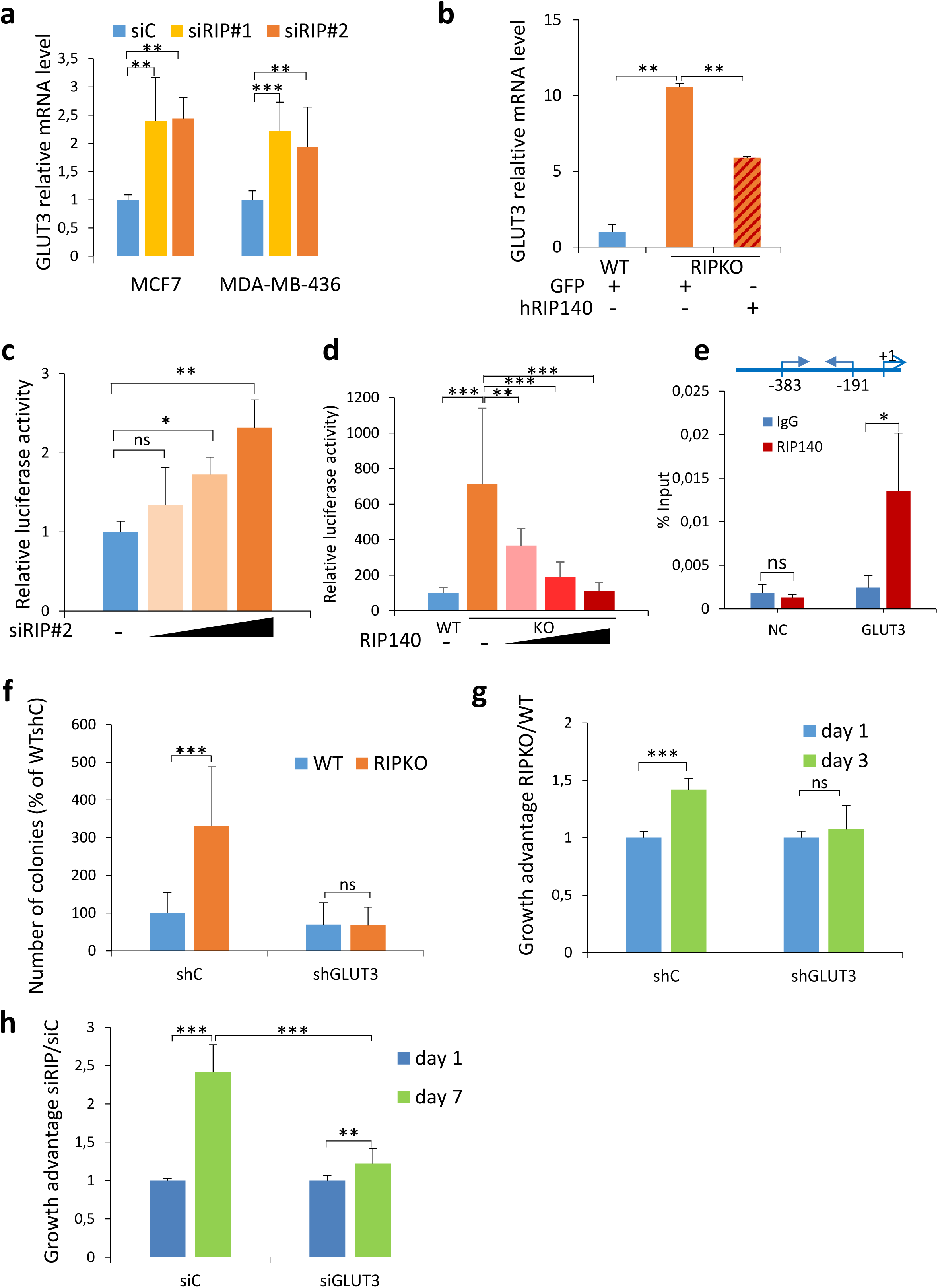
GLUT3 is essential for the growth advantage induced by RIP140-deficiency. **a**, *GLUT3* mRNA expression relative to 28S in MCF7 and MDA-MB-436 cells transfected with control siRNA (siC) or two different RIP140 siRNA (siRIP#1, siRIP#2). Values are normalized to control siRNA samples (mean ± SD, n=3, **p <0.01, ***p<0.001). **b**, *GLUT3* mRNA expression relative to RS9 in MEF #1 cells stably overexpressing a pEGFP plasmid (WT+GFP) and (RIPKO+GFP) or a pEGFP-hRIP140 plasmid (RIPKO+hRIP140). Values are normalized to WT+GFP samples (mean ± SD, n=3, **p <0.01). **c**, Luciferase activity assay in MDA-MB-436 cells transfected with RIP140 siRNA increasing doses, the luciferase reporter containing the promoter region of the GLUT3 gene and the luciferase reporter TK-Renilla. Luciferase values were normalized to the Renilla luciferase control. The data were normalized to that of the control siRNA (mean ± SD, n=3, *p<0.05, **p<0.01, *ns* not significant). **d**, Luciferase activity assay in MEF #1 cells transfected with increasing doses of pef-cMyc-RIP140 vector, the luciferase reporter containing the promoter region of the GLUT3 gene and the luciferase reporter TK-Renilla. Luciferase values were normalized to the Renilla luciferase control and to the WT samples (mean ± SD, n=3, ***p<0.001). **e**, ChIP assays were performed with an anti-RIP140 antibody or an irrelevant antibody (IgG) to determine the recruitment of RIP140 to the GLUT3 promoter. The GLUT3 promoter and the specific primers used in ChIP-qPCR are described (+1=transcription starting site). NC: non coding sequence. Enrichments are represented as percentages of input (mean ± SD, n=3, *p <0.05). **f**, Number of colonies from a colony formation in soft agar assay of transformed MEF #1 transfected with shRNA control or GLUT3 expression lentivirus. The data are expressed as percentages of WT shControl samples (shC) (mean ± SD, n=3, ***p <0.001, *ns* not significant). **g**, 3-(4,5-dimethylthiazol-2-yl)-2,5-diphenyltetrazolium Bromide (MTT) assay in MEF #1 transfected with shRNA control or GLUT3 expression lentivirus. The data are normalized to day 1 values for each shRNA and to WT samples representing the growth advantage of RIPKO MEFs over WT MEFs (mean ± SD, n=3, ***p <0.001, *ns* not significant). **h**, Cell viability assessed by crystal violet staining of MDA-MB-436 cells transfected with RIP140 siRNA (siRIP) combined with GLUT3 siRNA. The data are normalized to day 1 values for each siRNA and to control siRNA (siC) samples representing the growth advantage of siRIP140 transfected cells over siControl cells (mean ± SD, n=3, **p <0.01, ***p <0.001).

To investigate the mechanism by which RIP140 inhibits *GLUT3* expression, we first showed by luciferase reporter assays using the *GLUT3* promoter that RIP140 knock-down increased the luciferase activity of this reporter in MDA-MB-436 and MEFs cells (Fig. 4c, d) and that RIP140 overexpression decreased it in MEFs cells (Fig. 4d).

By chromatin immunoprecipitation (ChIP) assay in MDA-MB-436 cells, we observed the recruitment of RIP140 to the *GLUT3* promoter, indicating that the regulation by RIP140 occurred directly at the transcriptional level (Fig. 4e). Moreover, by analyzing ChIP-seq data in MCF7 cells [24], we found RIP140 bound in the vicinity of the *GLUT3 (SLC2A3)* gene. Altogether, these data identify the *GLUT3* gene as a transcriptional target of the trans-repressive activity exerted by RIP140.

To evaluate to which extent RIP140-deficient cells needed GLUT3 to grow, we first validated the induction of GLUT3 at the protein level in MEFs (Supplementary Fig. 4d) and then knocked it down with small hairpin RNA (shRNA) in MEFs, thus confirming the specifity of the GLUT3 antibody (Supplementary Fig. 4e). Knocking down GLUT3 abrogated the growth advantage induced by RIP140-deficiency as shown by either colony formation (Fig. 4f) or MTT (Fig. 4g) assays in MEFs. In MDA-MB-436 breast cancer cells, *GLUT3* silencing by siRNA abolished also the growth advantage provoked by RIP140 silencing (Fig. 4h and Supplementary Fig. 4f). Altogether, these data show that the increase of cell proliferation upon RIP140-deficiency requires *GLUT3* induction.

### RIP140 and p53 inhibit the expression of GLUT3 induced by HIF-2α

One direct DNA binding factor that could likely mediate RIP140 activity is p53, as it is a well-known inhibitor of the Warburg effect (Gomes et al., 2018). Thus, we investigated whether p53 mediated any of the transcriptional activities of RIP140 on *GLUT3* expression. Using a luciferase reporter assay, we showed that increasing doses of p53 inhibited the luciferase activity driven by the *GLUT3* promoter (Fig. 5a), and that RIP140 reinforced this inhibition (Fig 5b). Of interest, RIP140 deficiency resulted in decreased p53 expression at the protein and mRNA levels (Fig. 5c and Supplementary Fig. 5a). Therefore, RIP140-deficient cells exhibit many of the characteristics of cells under hypoxia such as cell proliferation, increased glycolysis and inactivation of p53 [12].

**Fig. 5.**
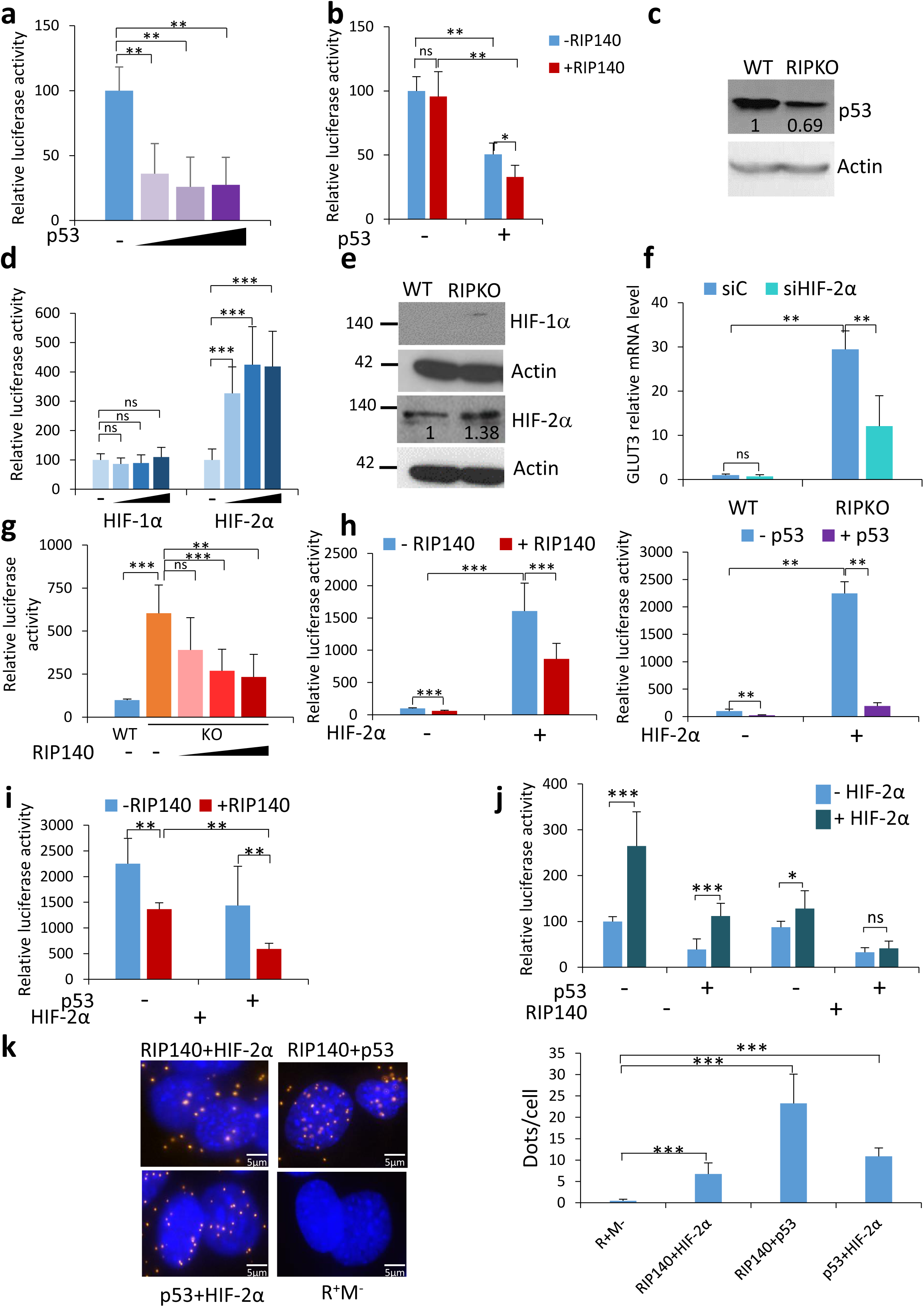
RIP140 and p53 inhibit the expression of GLUT3 induced by HIF-2α. **a**, Luciferase activity assay in MEF #1 RIPKO cells transfected with increasing doses of pcDNA3.1-p53 vector, the luciferase reporter containing the promoter region of the GLUT3 gene and the luciferase reporter TK-Renilla. Luciferase values were normalized to the Renilla luciferase control and to the values of samples without p53 (mean ± SD, n=4, **p <0.01). **b**, Luciferase activity assay in MEF #1 RIPKO cells transfected with pef-cMyc-RIP140, pcDNA3.1-p53 or empty vectors, the luciferase reporter containing the promoter region of the GLUT3 gene and the luciferase reporter TK-Renilla. Luciferase values were normalized to the Renilla luciferase control and to the values of samples transfected with empty vectors (mean ± SD, n=3, **p <0.01, *ns* not significant). **c**, The levels of p53 and Actin were assessed by western blot in MEF #1. A representative blot is shown with densitometry analysis. The intensity of p53 bands are normalized to that of Actin and to the WT sample. **d**, Luciferase activity assay in MEF #1 RIPKO cells transfected with pcDNA3.1-HIF-1α or pcDNA3.1-HIF-2α, the luciferase reporter containing the promoter region of the GLUT3 gene and the luciferase reporter TK-Renilla. Luciferase values were normalized to the Renilla luciferase control and to the values of samples transfected with empty vectors (mean ± SD, n=3, ***p <0.001, *ns* not significant). **e**, The levels of HIF-1α, HIF-2α and Actin were assessed by western blot in MEF #1 cells. Representative blots are shown with densitometry analysis for HIF-2α. The intensity of HIF-1α and HIF-2α bands are normalized to that of Actin and to the WT samples. **f**, Glut3 mRNA expression relative to RS9 in MEF #1 cells transfected with control siRNA (siC) or HIF-2α siRNA (siHIF-2α). Values are normalized to control siRNA WT samples (mean ± SD, n=3, **p < 0.01, *ns* not significant). **g**, Luciferase activity assay in MEF #1 cells transfected with increasing doses of pef-cMyc-RIP140 vector, the luciferase reporter containing hypoxia inducible response elements and the luciferase reporter TK-Renilla. Luciferase values were normalized to the Renilla luciferase control and to the WT samples (mean ± SD, n=3, ***p <0.001, *ns* not significant). **h**, Luciferase activity assay in MEF #1 RIPKO cells transfected with pcDNA3.1-HIF-2α, the luciferase reporter containing hypoxia inducible response element and the luciferase reporter TK-Renilla and pef-cMyc-RIP140 vector (left panel) or pcDNA3.1-p53 (right panel). Luciferase values were normalized to the Renilla luciferase control and to the values of samples transfected with empty vectors (mean ± SD, n=4, **p <0.01, ***p <0.001). **i**, Luciferase activity assay in MEF #1 RIPKO cells transfected with pcDNA3.1-HIF-2α plasmid, the luciferase reporter containing hypoxia inducible response elements and the luciferase reporter TK-Renilla, pef-cMyc-RIP140 and pcDNA3.1-p53 vectors. Luciferase values were normalized to the Renilla luciferase control and to the values of samples transfected with empty vectors (mean ± SD, n=3, **p <0.01). **j**, Luciferase activity assay in MEF #1 RIPKO cells transfected with pcDNA3.1-HIF-2α plasmid, the luciferase reporter containing the promoter region of the GLUT3 gene and the luciferase reporter TK-Renilla, pef-cMyc-RIP140 and pcDNA3.1-p53 vectors. Luciferase values were normalized to the Renilla luciferase control and to the values of samples transfected with empty vectors (mean ± SD, n=3, ** p <0.05, ***p <0.001, *ns* not significant). **k**, *Left panel*: In situ proximity ligation assay (PLA) was performed in fixed MEF #1 WT cells to visualize interaction between endogenous RIP140, p53 and HIF-2α. Nuclei were counterstained with Hoechst 33342 and negative controls were performed by incubating fixed cells without the primary antibodies (R+M-). *Right panel*: 5 different fields were randomly chosen for each condition in three independent experiments. Quantification of PLA is represented as mean±SD of the number of dots per cell (n=3, ***p <0.001). Scale bar, 5 µM.

To investigate the role of hypoxia in *Glut3* overexpression upon RIP140-deficiency, we first up-regulated the HIFs with a luciferase gene reporter driven by the *GLUT3* promoter. We found that the overexpression of HIF-2α, but not HIF-1α, increased the activity of the *GLUT3* promoter (Fig. 5d). Indeed, HIF-1α regulatory regions on *GLUT3* gene are not located within the promoter region [30]. We also found that HIF-2α, and not HIF-1α, was overexpressed in RIP140-deficient cells (Fig. 5e and Supplementary Fig. 5b). Interestingly, HIF-2α silencing reduced the expression of *Glut3*, only in RIP140 KO MEFs, suggesting that HIF-2α is responsible, at least in part, for *Glut3* overexpression upon RIP140 deficiency (Fig. 5f and Supplementary Fig. 5b).

To evaluate whether RIP140 could target HIFs activity, we first measured the effect of RIP140 on the transcriptional activity of HIFs using a luciferase reporter assay. We found that the transcriptional activity of a reporter gene containing hypoxia-response elements was higher in RIPKO than in WT MEFs and that increasing doses of RIP140 repressed the basic activity of this reporter gene (Fig. 5g). p53 is known to inhibit the transcriptional activity of HIFs (Blagosklonny et al., 1998). We then wondered whether RIP140 and p53 could cooperate to inhibit HIF transcriptional activity. We first looked at the effect of RIP140 or p53 individually on the transcriptional activity of the reporter gene containing hypoxia-response elements when transactivated by HIF-1α or HIF-2α. RIP140 or p53 were both capable of inhibiting HIF-α transcriptional activity separately (Fig. 5h and Supplementary Fig. 5c). Of interest, the repressive activity was stronger when RIP140 and p53 were expressed together, suggesting that RIP140 and p53 cooperate to inhibit HIF-α transcriptional activity (Fig. 5i and Supplementary Fig. 5d). The same synergistic repressive effect was observed on the *GLUT3* promoter when transactivated by HIF-2α (Fig. 5j).

Performing a proximity ligation assay demonstrated that HIF-2α, p53 and RIP140 interacted with each other (Fig. 5k). Altogether, these data suggested that the molecular mechanisms by which RIP140 inhibits *GLUT3* transcription relies on the repression of HIF transactivation through the formation of a ternary complex with p53.

### The prognostic value of RIP140 is correlated with the levels of GLUT3 expression

Our experimental data demonstrated that RIP140 inhibits efficiently GLUT3 expression in synergy with p53, by antagonizing HIF-2α function, leading to a decrease in cancer cell proliferation. We therefore hypothesized that, in breast cancers, low GLUT3 levels could be a marker of such anti-tumor activity of RIP140 and therefore, that RIP140 might be associated with an increased overall survival for patients with tumor expressing a reduced level of GLUT3, as a surrogate of RIP140 anti-tumor activity.

Using Cox proportional hazard regression [32], we analyzed RNAseq data obtained from 1068 breast tumor samples from the TCGA dataset as described previously [33] (Fig. 6). We first checked the prognostic values of the expression of each gene by analyzing patient overall survival at 60 months. High RIP140 (NRIP1) expression was correlated with good prognosis whereas high GLUT3 (SLC2A3) was associated with bad prognosis (Fig. 6a). Then we used the median as a cutoff value for classification of patients into two groups of tumors with low and high GLUT3 expression, respectively. Using Kaplan-Meier plots, we investigated whether RIP140 expression correlated with OS at 60 months within both groups using the same cutoff values. Interestingly, and as expected, RIP140 expression was significantly associated with increased overall survival in low GLUT3 expression group but not in high GLUT3 expression group (Fig. 6b).

**Fig. 6.**
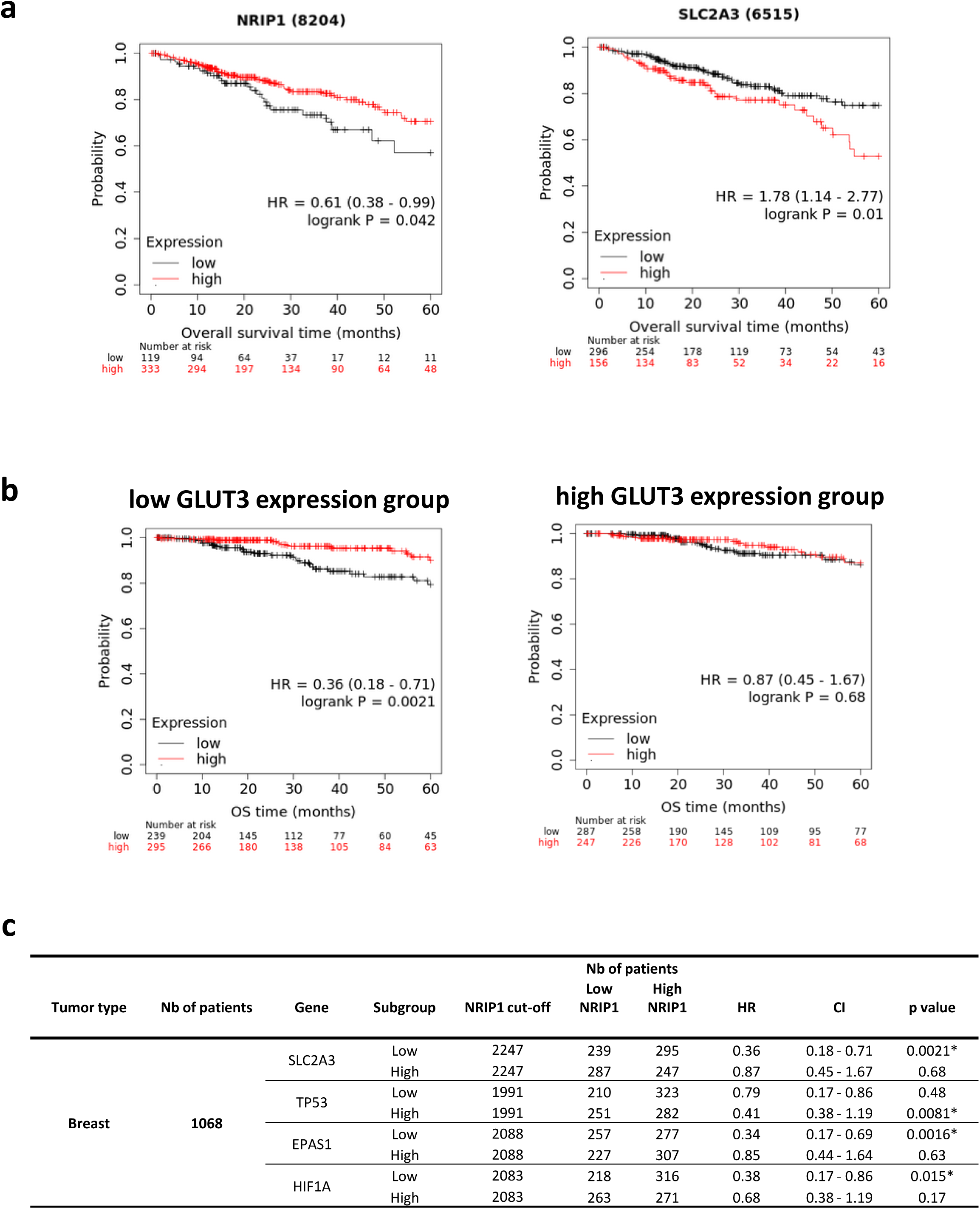
The prognostic value of RIP140 is correlated with the levels of GLUT3 expression. Kaplan-Meier analysis plotting the survival curve of 1068 cases of breast cancer with statistical significance being assessed using the log-rank test from the TCGA dataset. **a**, Kaplan-Meier curves showed worse overall survival rates for breast cancer patients with low RIP140 (NRIP1) expression compared to patients with high RIP140 expression (P=0.042; left panel). Kaplan-Meier curves showed worse overall survival rates for breast cancer patients with high GLUT3 (SLC2A3) expression compared to patients with low GLUT3 expression (P=0.01; right panel). **b**, The median of GLUT3 expression was used to define patients into two groups; the low GLUT3 expression group (left panel) and the high GLUT3 expression group (right panel). Kaplan-Meier curves showed better overall survival rates for breast cancer patients with high RIP140 expression in the low GLUT3 expression group (P=0.0021; left panel). On the contrary, RIP140 prognostic value was not significant in the high GLUT3 expression group (P=0.68; right panel). **c**, Groups have been defined on the basis of the median p53 (TP53), HIF-2α (EPAS1) and HIF-1α (HIF1A) expression. RIP140 prognostic value was significant in high p53 (P=0.0081), low HIF-2α (P=0.0016) and low HIF-1α (P=0.015).

We pursued our hypothesis by analyzing the prognostic value of RIP140 based on p53 (TP53), HIF-2α (EPAS1) and HIF-1α (HIF1A) expression levels. Patients were stratified again into low and high expression groups by using, as cutoff values, the median of either TP53, HIF-2α, or HIF-1α expression (Fig. 6c and Supplementary Fig. 6a). Strikingly, RIP140 expression was significantly associated with good prognosis in high p53 and low HIF-2α or HIF-1α expression groups, confirming the mechanism of RIP140 anti-tumor activity provided by our experimental data (Fig. 6c). Moreover, the prognostic value of RIP140 was significantly associated with good prognosis in low GLUT3 expression group but not in high GLUT3 expression group in colon and stomach cancers, suggesting that a reduced level of GLUT3 could reflect RIP140 anti-tumor activity in other types of cancer (Supplementary Fig. 6b).

## Discussion

Glycolysis is essential for supporting the rapid proliferation of tumor cells. Our data reveal that the transcription coregulator RIP140 inhibits the glycolysis-dependent proliferation of breast cancer cells by impeding glycolysis through the blockade of *GLUT3* expression *via* a mechanism involving a p53-mediated inhibition of HIF activation.

Our results first demonstrate that the *glucose deprivation or glycolysis blockade abolished the gain in cell proliferation caused by the decrease or loss of RIP140 expression*, showing the glucose dependency of RIP140-deficient cells. Furthermore, our data demonstrate that *down-regulating Glut3 expression provoked the same inhibition of RIP140-deficient cell proliferation*. H-Ras transformation is well recognised to drive cancer cells towards glycolysis and glycolysis is essential for H-Ras transformation [6]. Of note, transformed RIPKO cells were more sensitive to glucose starvation than immortalized RIPKOMEFs (Fig. 2b and Supplementary Fig. 2a), suggesting that RIP140-deficiency might influence the transformation process by regulating glycolysis or that Ras transformation could enforce the potential of RIP140. However, RIP140 potential was also observed in Ras-independent cells, such as immortalized MEFs or the human cancer line MDA-MB-436.

RIP140 potential was also independent of PTEN which is another driver of cancer glycolysis. Indeed, we observed the same effect of RIP140-deficiency in the PTEN-competent breast cancer cell line MCF7 and in the PTEN-negative MDA-MB-436 breast cancer cell line.

Interestingly, RIP140 does not regulate the metabolism of glutamine although this amino acid is also important for the proliferation of cancer cells (Jin et al., 2016). Indeed, the proliferation of WT and RIP140 KO cells was equally impacted by glutamine deprivation (data not shown), suggesting that RIP140 specifically affects glycolysis.

Our tailored experiments describe the *mechanism by which RIP140 inhibits GLUT3 expression, which relies on the cooperation of RIP140 and p53* to inhibit the expression of *GLUT3* induced by HIF-2α (Fig. 5g). p53 wild-type seems to be dispensable for the cooperation since RIP140 silencing induced GLUT3 expression in p53 mutated cells such as the MDA-MB-436 breast cancer cell line. p53 could act as a facilitator of the RIP140 repressive effect. Proximity ligation assays allowed us to visualize the three partners in close proximity two-by-two, suggesting that they are involved in a ternary complex (Fig. 5h). Our data reveal for the first time that RIP140 interacts with p53 and HIF. Whether there is a ternary complex or if the interactions follow a temporal order will need to be defined in future studies. It is tempting to speculate that RIP140 could act as an integrator protein such as CBP/p300 as p53 and HIF antagonism relies on competition for p300 (Schmid et al., 2004). RIP140 could enter into the competition with CBP as it does for nuclear receptors (Chen et al., 2004). The interplay between p53 and HIFs remains a complex question and it is still debated whether their reciprocal influence has any direct consequences for metabolism in cancer. Our results add RIP140 as a new major player in this interplay, and provide an additional bond between cell metabolism and cancer progression.

The overexpression of *GLUT3* has been described in many types of cancer with poor outcomes (Benito et al., 2017). In breast cancer patients, we found that *RIP140 is associated with an increased overall survival of patients with tumor expressing a reduced level of GLUT3*, as a surrogate of RIP140 anti-tumor activity (Fig. 6). Cellular metabolism in cancer is currently being targeted in clinical trials with some success (Sborov et al., 2015). Our results suggest that these genes could be used as a signature to identify patients that could benefit of therapies targeting glycolysis and/or based on HIF inhibitors.

Altogether, our results enable us to propose a new model (Figure 7) explaining the transcriptional control of glycolysis-dependent cancer cell proliferation by a nuclear interplay between three actors that might be clinically relevant for breast cancer patients.

**Fig. 7.**
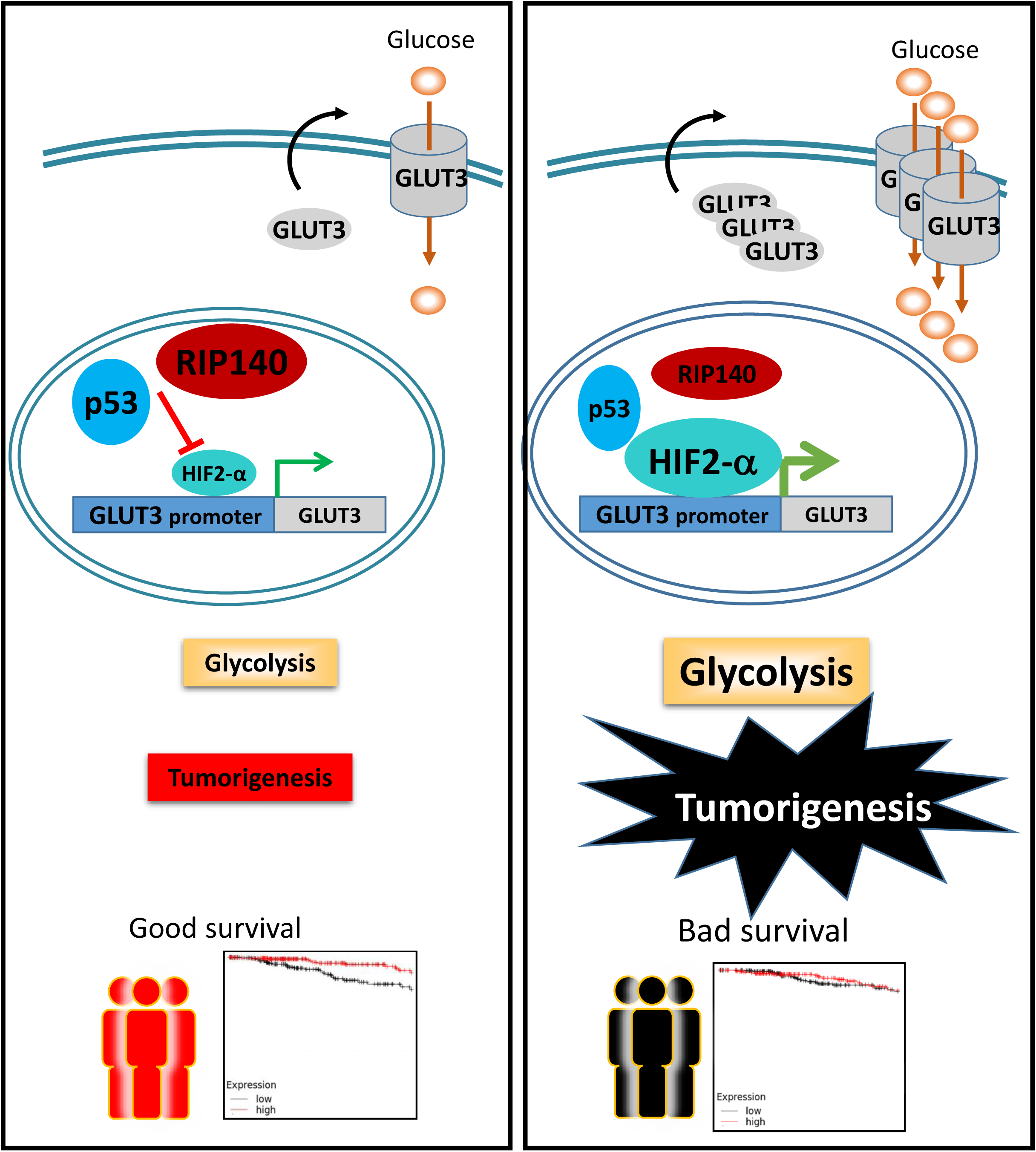
Model: RIP140 inhibits tumorigenesis by affecting glycolysis through the blockade of GLUT3 expression. RIP140 and p53 cooperate to inhibit the expression of GLUT3 induced by HIF-α. Glycolysis-dependent proliferation of breast cancer cells is reduced, due to a decrease in glycolysis. The prognostic value of RIP140 is associated with good survival in patients with low GLUT3, high p53 and low HIF-α (right panel). In patients with high HIF-α, low p53 and high GLUT3, RIP140 and p53 do not inhibit the transcriptional activity of HIF-α; GLUT3 is highly expressed and glycolysis is enhanced. In this sub-group, RIP140 expression level is not correlated with good survival. Double blue lines represent cell membranes. The grey square represents GLUT3 gene, the grey ovoid forms represent GLUT3 protein. The orange circle represents glucose.

## Materials and Methods

### Plasmids and reagents

RIP140-expressing vectors (pEFcmyc-RIP140 [40], pEGFP-RIP140[41]) and control vectors, pEGFP (Clontech), are described elsewhere. HA-HIF1alpha-pcDNA3 (Addgene plasmid # 18949) and HA-HIF2alpha-pcDNA3 (Addgene plasmid # 18950) plasmids were gifts from Dr Kaelin[42]. The GLUT3-Luc reporter gene was a gifts from Dr. Yuan (Liu et al., 2014).. 2-Deoxy-D-glucose (2DG, D6134)), 6-Aminonicotinamide (6AN; # A68203), 3-Bromopyruvate (BrP; #16490), MTT ((3-(4,5-dimethylthiazol-2-yl)-2,5-diphenyl tetrazolium bromide) (98%, CAS 298-93-1), Deferoxamine mesylate salt (D9533), Di(N-succinimidyl) glutarate (80424), crystal violet (C0775), anti-Mouse IgG-FITC antibody (F6257), monoclonal Anti-ß-Actin-Peroxidase antibody (A3854) were purchased from Sigma-Aldrich (Merck, Darmstadt, Germany). Puromycin (ant-pr-1) was purchased from Invivogen (France). The ATP Determination Kit (10700345) was from Fisher Scientific (France). Deoxy-D-glucose, 2-[1,2-3H (N)] (NET328A250UC) was from Perkin Elmer (France). GLUT3 shRNA (m) (sc-41219-V) were purchased from Santa Cruz Biotechnology (Dallas, USA). Rabbit polyclonal to RIP140 (ab42126) was from Abcam (Cambridge, UK). BrdU Hu-purified-clone B44 (#347580) was from Becton Dickinson (France). Ambion™ Silencer™ Pre-Designed siRNA specific of human GLUT3 (SLC2A3) was purchased from Fisher Scientific (#10446914, Illkirch, France)

#### Cell culture

Immortalized and transformed mouse embryonic fibroblasts (MEFs) were cultured in F12/Dulbecco’s modified Eagle’s medium supplemented with 10% fetal calf serum, 1% Penicillin/Streptomycin, 1mM Sodium Pyruvate and 10mM HEPES. Primary MEFs were prepared from 13.5 day wild-type (WT) or RIPKO mouse embryos [44] and genotyped by PCR. Immortalized MEFs were obtained by sequential passage, according to the 3T3 protocol [45], or by infection with retrovirus expressing SV40. After immortalization, MEFs were transformed with retrovirus expressing H-RasV12. Virus production, infection and transfection were performed as previously described[46]. Stably infected cells were selected with Puromycin (2.5µg/ml) after SV40 virus infection and Hygromycin (65µg/ml) after H-RasV12 virus infection. Four independently derived RIP140 WT and RIPKO MEF cell lines were established. MEF#1 were immortalized by the 3T3 protocol. MEF#2, #3 and #4 were transformed by the infection of SV40/H-RasV12 expressing retrovirus. Immortalized MEF#1 were transduced with lentiviral particles expressing shRNA against murine GLUT3 (sc-41219-V) according to the manufacturer’s protocol (Santa Cruz Biotechnology,). MEFs transfected with pEGFP or pEGFP-hRIP140 were cultured in the presence of 3.2µg/ml Puromycin, as the selection agent to establish stable cell lines. MCF7, PC3 and DU145 were cultured in F12/Dulbecco’s modified Eagle’s medium supplemented with 10% fetal calf serum, 1% Penicillin/Streptomycin. MDA-M-B436 and RKO were cultured in Dulbecco’s Modified Eagle Medium (DMEM) + GlutaMax (Thermo Fisher Scientific; Waltham, MA), supplemented with 10% FBS and 1% Penicillin/Streptomycin.

#### Real-time qPCR

Real-time qPCR was conducted as previously described [28]. Briefly, total RNA was extracted from cells using the Quick RNA^TM^ Miniprep kit (Zymo Research, Irvine, CA, USA) according to the manufacturer’s instructions. cDNA was synthetized with 1 µg of RNA and the qScript cDNA master mix (Quanta Bio, Beverly, MA, USA). mRNA expression was determined with a quantitative real time PCR SYBR Green SensiFAST™ SYBR® No-ROX Kit (Bioline, London, UK) on a Light Cycler® 480 Instrument II (Roche Life Sciences). Relative expression levels for the mRNAs of interest were normalized to 28S or RS9 housekeeping genes. See Supplemental Table 1 for the list of primer sequences.

### Protein detection

Protein expression was quantified by western blot analysis using the following antibodies: anti-cyclin A (1:500 sc-751; Santa Cruz Biotechnology, Dallas, TX, USA), anti-Histone H3 (phospho Serine10) (1:2,000 ab14955; Abcam Cambridge, UK), anti-GLUT3 (1:10000 ab191071; Abcam), anti-p53 (1:1000 1C12; Cell Signaling Technology, Leiden, The Netherlands), anti-HIF-1α (1:1000 NB100-105; Novus Biologicals, LLC (Centennial, USA)), anti-HIF-2α/EPAS1 (1:1000 NB100-122; Novus Biologicals, LLC (Centennial, USA)), and anti-Actin-HRP (1:10000 A3854, Merck, Darmstadt, Germany). Densitometry analysis were performed by using Image J software.

Immunofluorescence and proximity ligation assays (PLA) were performed as described[28].

#### Luciferase assay

For gene reporter assays, cells were plated in 96-well plates and transfected with JetPEI (Polyplus, Illkirch-Graffenstaden, France) according to the manufacturer’s protocol. Data were expressed as relative firefly luciferase activity normalized to renilla activity. For all experiments, data were collected from at least three biological replicates.

#### Chromatin immunoprecipitation

Approximately one hundred and twenty million cells were harvested per experiment. Briefly, protein–DNA complexes were first cross-linked, using 2mM Di (N-succinimidyl) Glutarate (DSG), for 50 min on a rotor, then the procedure was performed using High-Sentivity kit (Active Motif, Shanghai) and according to manufacturer’s protocol. Chromatin was sonicated for thirty seconds on ice followed by thirty seconds off, for a total of thirty minutes. Immunoprecipitations were performed with 30µg of protein-DNA complexes and 4µg of RIP140 antibody (Ab42126; Abcam). The purified DNAs were amplified with SYBR Green SensiFAST™ SYBR® No-ROX Kit (Bioline, London, UK) by real-time qPCR.

### Cell growth analysis

#### MTT assay

Cell proliferation assays were performed using MTT as previously described [28]. Data were normalized to the absorbance value at day 1. Experiments were performed three times with at least sextuplets. For glucose starvation experiments, cells grew for 24 hours in complete media. Then cells were washed twice with PBS and starvation medium (DMEM5030 supplemented with 10% dialyzed FBS, 2mM Glutamine, 1% Penicillin/Streptomycin, HEPES 10mM, 1mM Sodium Pyruvate, 3.7g/l Sodium Bicarbonate) was added to the cells supplemented, or not, with indicated glucose concentrations.

#### xCELLigence analysis

The Real-time Cell Analyzer DP instrument (Agilent, Santa Clara, USA) was placed in a humidified incubator maintained at 37°C with 5% CO_2_. Cells were plated at 1500 cells/well into 16-well E-Plates. The impedance value of each well was automatically monitored every hour for up to the indicated times by the xCELLigence system. Cell proliferation is represented by an index that reflects changes in electrical impedance matching to cellular coverage of the electrode sensors.

#### Soft agar colony assay

Soft agar experiments were performed as described previously[46]. Briefly, transformed MEFs (8.10^4^ cells) were mixed with culture medium containing 0.4% agar and seeded into 6-well plates coated with 1% agar in F12/DMEM containing 10% FBS in at least triplicate. Medium was changed twice a week. After 20 days, colonies were stained with 0.5% crystal violet solution, photographed and counted with ImageJ software

#### Mouse xenograft assays

Transformed MEF (5. 10^5^ cells) were resuspended in 100 μl of RPMI medium (Thermo Fisher Scientific; Waltham, MA) and injected subcutaneously into the dorsal area of 4-6 week-old immuno-deficient athymic female mice (Charles River, Écully, France). After 10-14 days, tumor volumes were measured every 3-4 days with a caliper. Before reaching tumor volume limit point, fibrosarcomas were excised, photographed and weighed.

### Metabolic measurements

#### Glucose uptake

One day after cell seeding at 50,000 cells/well into 24-w plates, cells were washed once with PBS and medium was replaced with low-glucose medium (DMEM5030 supplemented with 10% dialyzed FBS (Thermo Fisher Scientific, Waltham, MA), 2mM Glutamine, 1% Penicilline/Streptomycin, HEPES 10mM, 1mM Sodium Pyruvate, 3.7g/l Sodium Bicarbonate, 0.25mM Glucose). Then 1µCi/ml of Deoxy-D-glucose, 2-[1,2-3H(N)] (Perkin Elmer, Waltham, MA) was added for 30 minutes at room temperature. Cells were washed once with PBS and then lysed with 200µl of 1%SDS; 10mM Tris pH 7.5. 100µl was used to count radioactivity with a Perkin Elmer liquid scintillation analyzer Tri-Carb 2900TR and 5µl was used to quantify proteins for normalization.

#### Cellular ATP measurement

ATP content was determined using the ATP determination kit according to the manufacturer’s instructions (Molecular Probes, Thermo Fisher Scientific,Waltham, MA). Cells were washed with iced-cold 1x PBS and extracted in an ATP-releasing buffer containing 100 mM potassium phosphate buffer at pH 7.8, 2 mM EDTA, 1 mM dithiothreitol, and 1% Triton X-100. Then, 5 μl of lysate was used for protein determination by the DC Protein Assay (Bio-Rad Laboratories, Hercules, CA).

#### Seahorse experiments

Extracellular acidification rate (measured in mpH/min) was monitored using an XFe24 extracellular flux analyzer from Seahorse Bioscience (Agilent, Santa Clara, CA) following the manufacturer’s protocol. Experiments were carried out on confluent monolayers. Briefly, cells were seeded 24 hours before experiments at a density of 35,000 cells/well (24-well). Before starting measurements, cells were washed once with PBS and medium was replaced with Seahorse XF Base Medium supplemented with 2mM glutamine without glucose at pH 7.4) and were placed into a 37 °C non-CO2 incubator for 1 hour prior to the assay. Glucose, oligomycin, and 2-DG were diluted into XF24 media and loaded into the accompanying cartridge to achieve final concentrations of 10 mM, 1 μm, and 100 mM, respectively. Injections of the drugs into the medium occurred at the time points specified. Each cycle consisted of 3 min mixing, 3 min waiting and 3 min measuring. Data were transformed with Agilent Seahorse Wave software to export glycolysis parameters. Values were expressed after normalization to the protein content of each well and to the values just before glucose addition.

#### Glucose and Lactate assays

Cells were seeded in culture dishes and cultured for 8 h. The culture medium was then changed and cells were incubated for an additional 16 h. Subsequently, the culture medium was collected for determination of glucose concentration and lactate levels using a Glucose assay kit (Amplite ^TM^ Glucose Quantitation Assay kit, AAT Bioquest, Sunnyvale, CA) and a Lactate assay kit (MAK064, Merck, Darmstadt, Germany) according to the manufacturer’s instructions. Glucose consumption was calculated as the difference in glucose concentration between fresh medium and cell supernatant. Lactate production was determined as the difference in lactate concentration between cell supernatant and fresh medium. Data were normalized to final cell counts and incubation time.

#### Survival analysis

RNA sequencing (RNA-seq) data were utilized from the Cancer Genome Atlas (TCGA) using the KM-plot Private Edition (http://kmplot.com) [33]. Using Cox proportional hazards regression, RNAseq data obtained from breast cancer samples were analyzed as previously described [32]. The Kaplan-Meier method was used to estimate overall survival calculated from diagnosis to death. Patients lost to follow-up were censored at the time of last contact.

#### Statistical analysis

Data are expressed as mean ± SD. Statistical analysis was conducted via StatEL (www.adscience.fr). The Mann–Whitney U-test was used to compare two independent groups. The p values less than 0.05 were considered to be statistically significant.

---

**Table 1.**
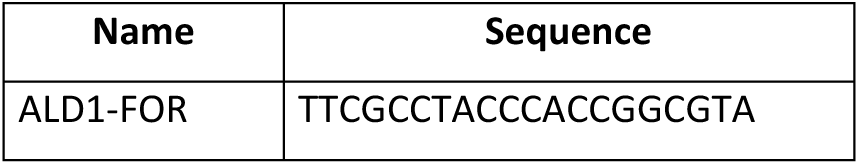
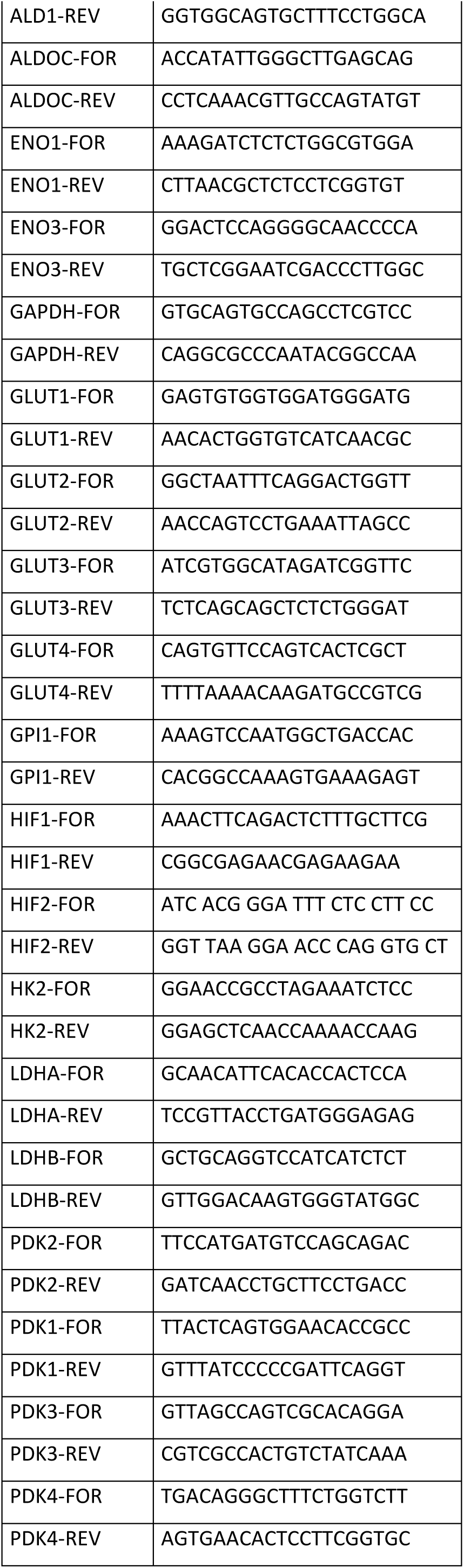
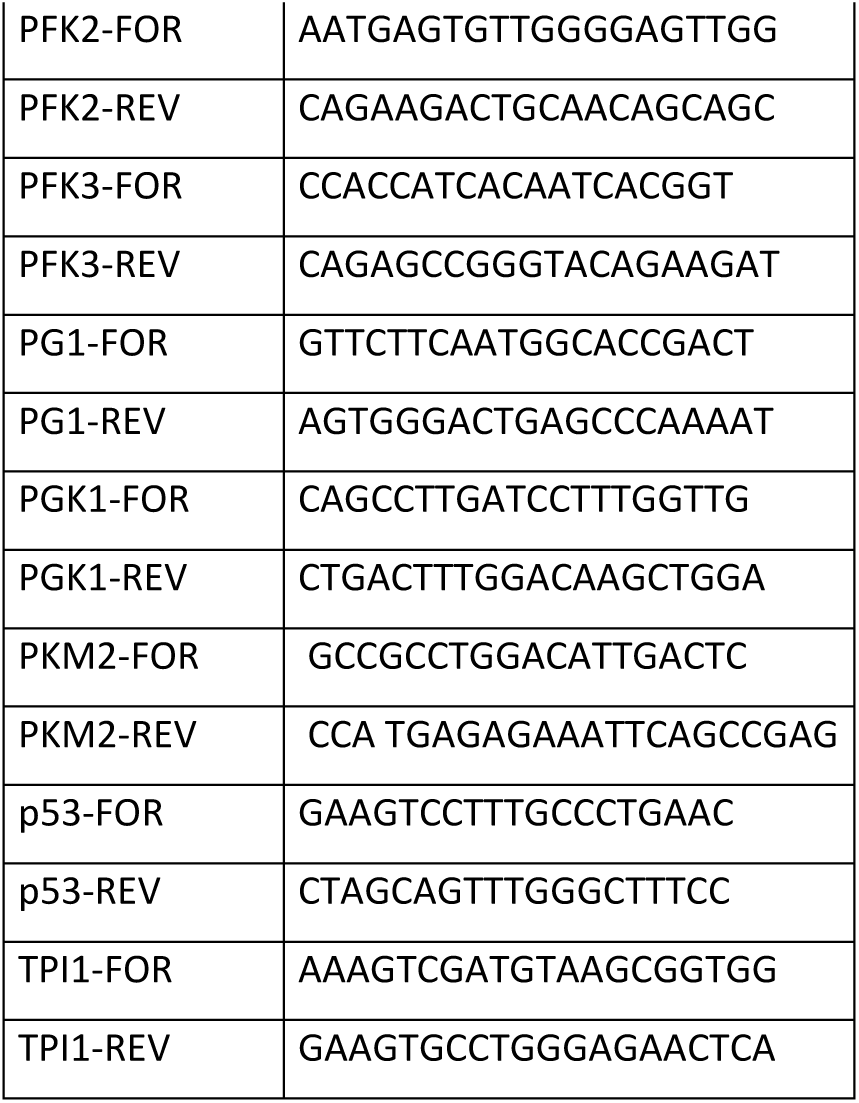
Murine primer sequences.

**Table 2.**
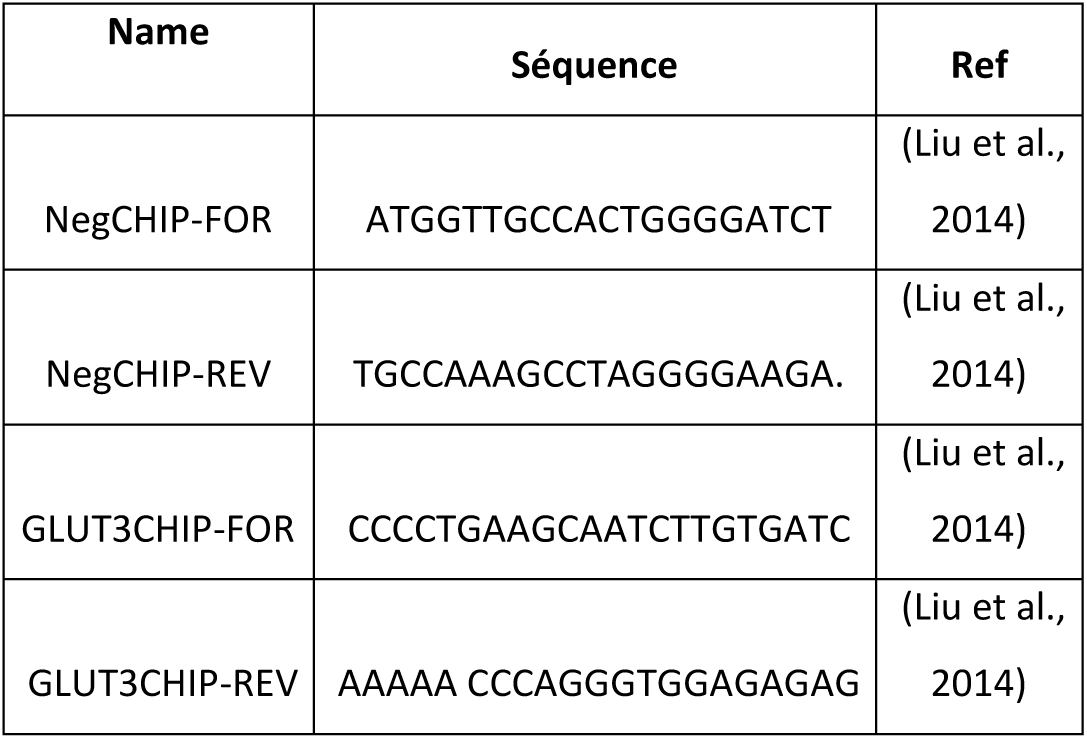
ChIP primer sequences.

**Table 3.**
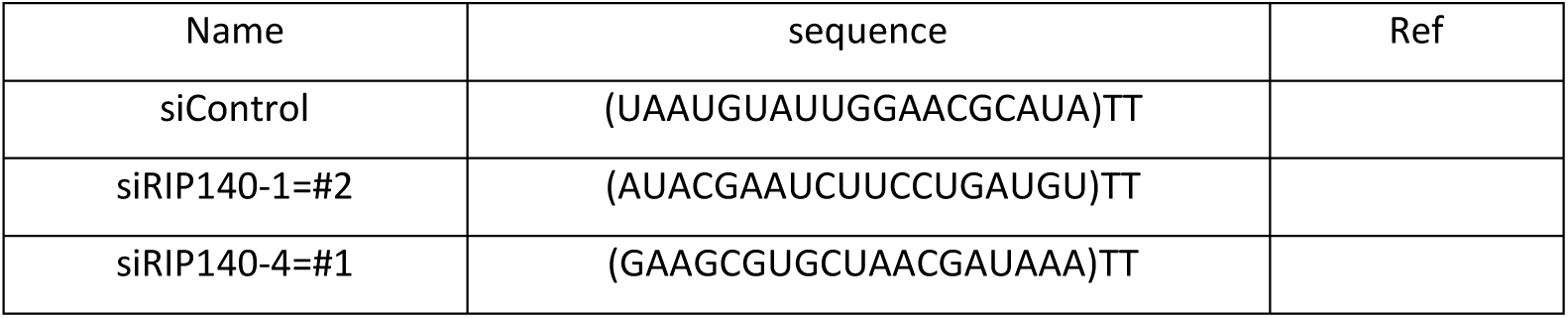
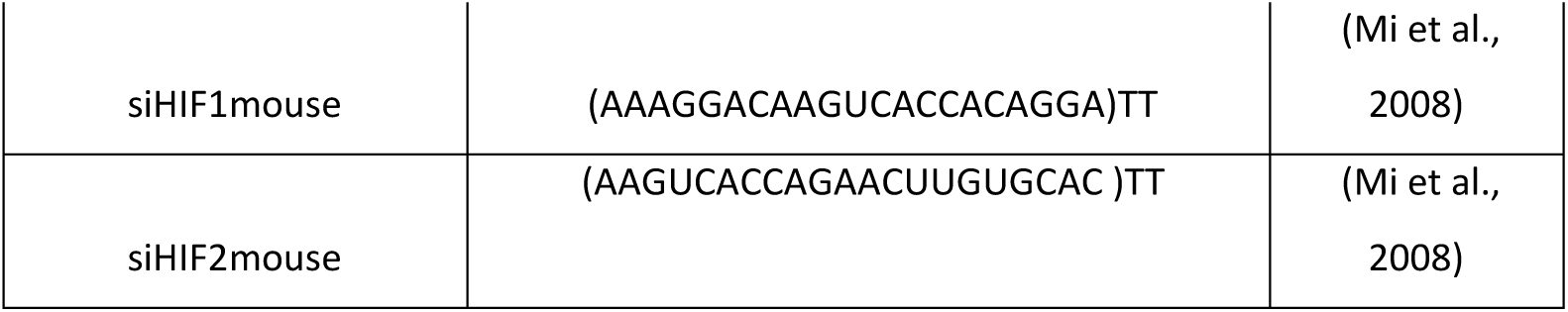
siRNA sequences.

## Statements and Declarations

### Author contributions

V.J., D.G., S.F., L.K.L., S.B., S.J. and C.T. performed the experiments. S.F. and L.K.L contributed to the *in vivo* experiments. S.F. generated MEF #2. V.J. and S.J. performed breast cancer cell proliferation experiments. V.J. performed ChIP experiments, glycolysis inhibition and siRNA experiments in cancer cells, and glucose starvation experiments. D.G performed immunoblots, ATP measurement, 2DG treatment experiments, cell proliferation assays in MEF#1. S.B. performed luciferase reporter assays. C.T. generated MEF #3 and #4, performed Seahorse, glucose uptake, xCELLigence, soft agar, PLA, shRNA experiments, luciferase reporter assays and immunoblots. V.J, V.C. and C.T. designed the studies and interpreted the data. C.T. wrote the manuscript. V.C. reviewed and approved the manuscript. All authors discussed and revised the manuscript. V.J. was supported by a fellowship from the French Ministry of Research.

### Data Availability

The links to publicly archived datasets analyzed in this study and all materials are available on request from the corresponding author.

### Ethical approval and consent to participate

All experiments on mice were performed in accordance with the French guidelines for experimental animal studies (agreement n° 201603101538202). No human samples were used in the study. No human research participants were used in the study.

## Acknowledgements

We thank Dr. Yuan, Dr. Karadimitris and Prof. Schymkowitz for the kind gift of plasmids, and Drs. Lapierre and Palassin for the generation of immortalized MEF #1 and stable MEF#1 overexpressing hRIP140, respectively. Thanks also to Dr. Stallcup for proofreading the manuscript.

## Supplementary Figure Legends

**Supplementary Fig. 1.**
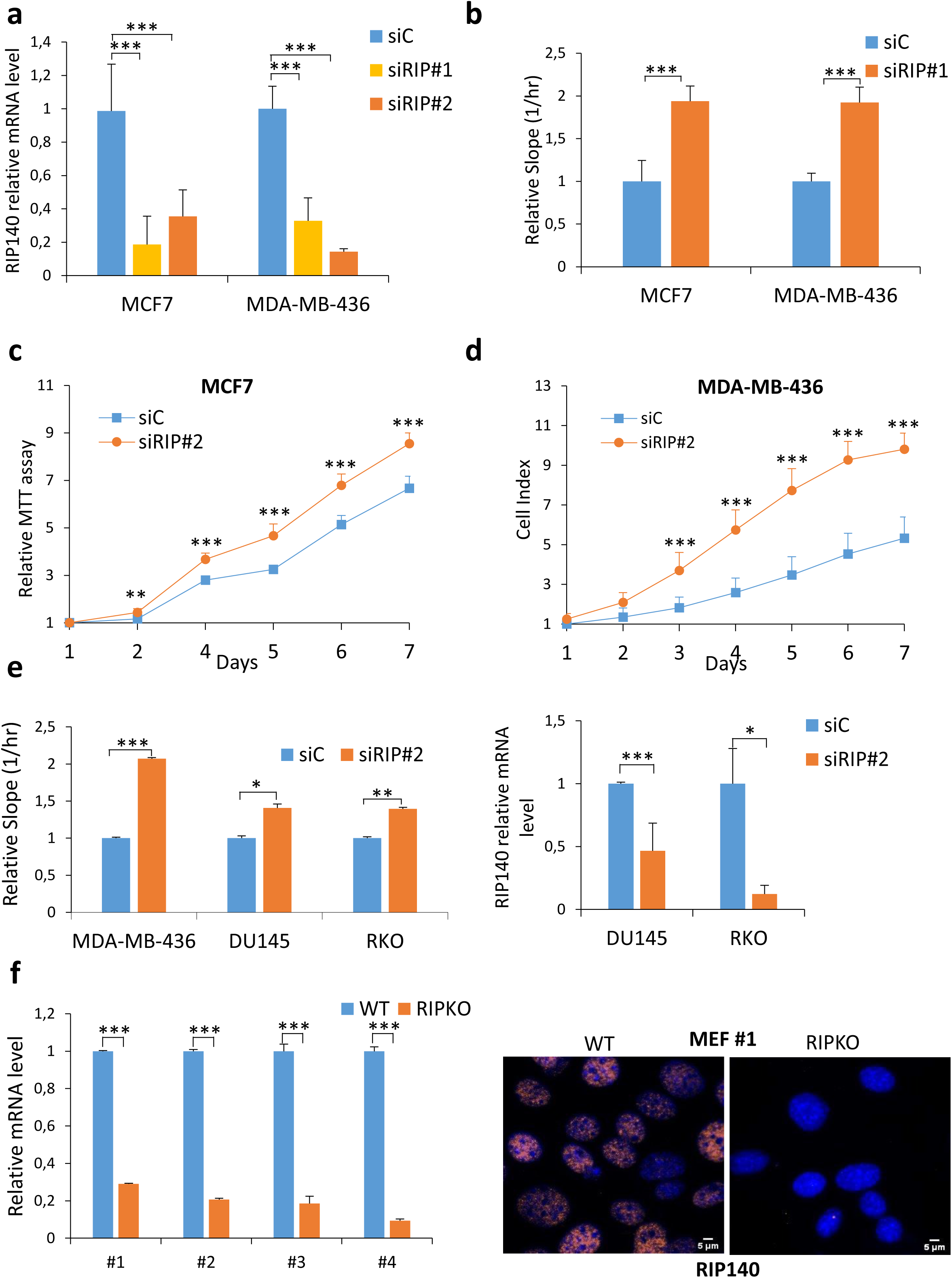
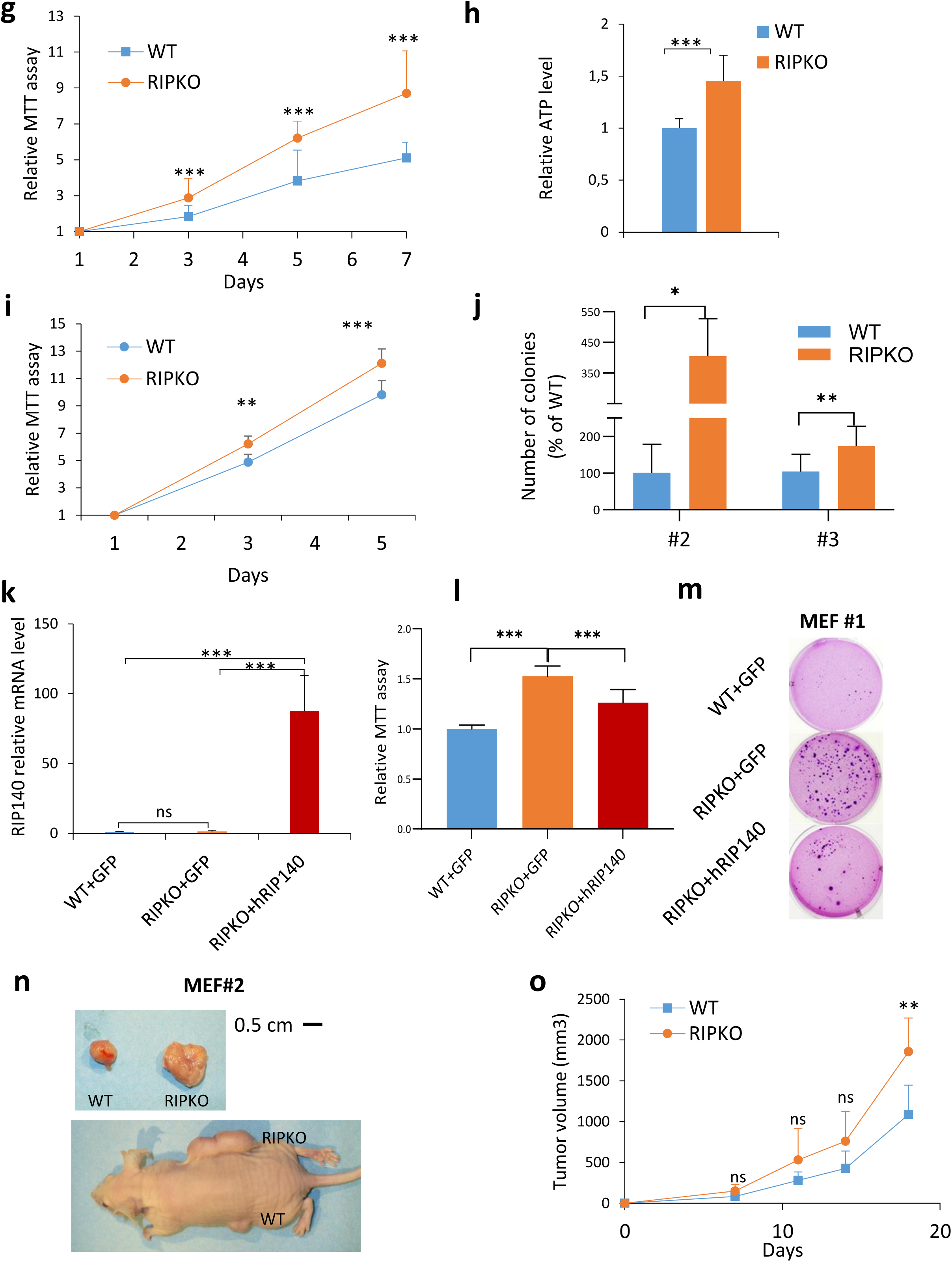
RIP140-deficiciency enhances cell proliferation and tumorigenesis. **a**, RIP140 mRNA expression relative to 28S in MCF7 and MDA-MB-436 cells transfected with control siRNA (siC) or RIP140 siRNAs (siRIP#1 used in Fig. 1a, b; siRIP#2 used in Supp Fig. 1c, d, e) (mean ± SD, n=3, ***p < 0.001). **b**, The slope of the curves was extracted using the RTCA Software from the curves in Fig. 1a and 1b (mean ± SD, n=3, ***p <0.001). **c**, 3-(4,5-dimethylthiazol-2-yl)-2,5-diphenyltetrazolium Bromide (MTT) assay in MCF7 cells transfected with control siRNA (siC) or RIP140 siRNA (siRIP#2). Values are normalized to day 1 (mean ± SD, n=3, ** p <0.01, ***p <0.001). **d**, Live measurements of cell proliferation were performed with the xCELLigence RTCA DP instrument in MDA-MB-436 cells transfected with control siRNA (siC) or RIP140 siRNA (siRIP#2) (mean ± SD, n=4, **p <0.01, ***p <0.001). **e**, *Left panel*: The slopes of the curves were extracted using the RTCA Software from Supplementary 1d for MDA-MB-436 and from live measurements of cell proliferation performed with the xCELLigence RTCA DP instrument in DU145 and RKO transfected with control siRNA (siC) or RIP140 siRNA (siRIP#2) (mean ± SD, n=3, *p <0.05, **p <0.01, ***p <0.001). *Right panel*: RIP140 mRNA expression relative to 28S in DU145 and RKO cells transfected with control siRNA (siC) or RIP140 siRNA (siRIP#2). (mean ± SD, n=3, *p <0.05, **p <0.01, ***p <0.001) **f**, RIP140 mRNA expression relative to RS9 in MEFs used in the study and generated from four different breedings. MEF#1 were immortalized by the 3T3 protocol. MEF#2, #3 and #4 were transformed by the infection of SV40/H-RasV12 expressing retrovirus (left panel, mean ± SD, n=6, ***p <0.001). Immunofluorescence imaging of RIP140 protein in MEF #1. Hoechst 33342 was used for nuclei detection. Scale bar: 5 μm (right panel). **g**, 3-(4,5-dimethylthiazol-2-yl)-2,5-diphenyltetrazolium Bromide (MTT) assay in MEF #1. Values are normalized to day 1 (mean ± SD, n=4, ***p <0.001). **h**, ATP level from MEF #1 was expressed in nM ATP per 10 μg protein units and then normalized with respect to the level in WT samples (mean ± SD, n=3, ***p <0.001). **i**, 3-(4,5-dimethylthiazol-2-yl)-2,5-diphenyltetrazolium Bromide (MTT) assay in SV40/H-RasV12-transformed RIP140 WT and KO MEF #3 (mean ± SD, n=3, ***p <0.001). **j**, Number of colonies from a colony formation in soft agar assay of transformed MEF #2 and SV40/H-RasV12-transformed MEF #3. Values are expressed as percent of WT samples (mean ± SD, n=3, *p <0.05, **p <0.01). **k**, RIP140 mRNA expression relative to RS9 in MEF #1 stably overexpressing a pEGFP plasmid (WT+GFP) and (RIPKO+GFP) or a pEGFP-hRIP140 plasmid (RIPKO+hRIP140). Primers used are specific for the human RIP140 sequence (mean ± SD, n=5, **p <0.01, *ns* not significant). **l**, 3-(4,5-dimethylthiazol-2-yl)-2,5-diphenyltetrazolium Bromide (MTT) assay in MEF #1 stably overexpressing a pEGFP plasmid (WT+GFP) and (RIPKO+GFP) or a pEGFP-hRIP140 plasmid (RIPKO+hRIP140) at day 6. Values are normalized to day 1 and to WT samples (mean ± SD, n=3, **p <0.01). **m**, Representative pictures of colony in soft agar assay of MEF #1 stably overexpressing a pEGFP plasmid (WT+GFP) and (RIPKO+GFP) or a pEGFP-hRIP140 plasmid (RIPKO+hRIP140) quantified in Fig. 1f. **n**, Pictures of xenografted nude mice with MEF #2 from Fig. 1e. **o**, Tumor growth curve of H-RasV12-MEF #1 xenografted in nude mice (mean ± SD, n=6, **p <0.01, *ns* not significant).

**Supplementary Fig. 2.**
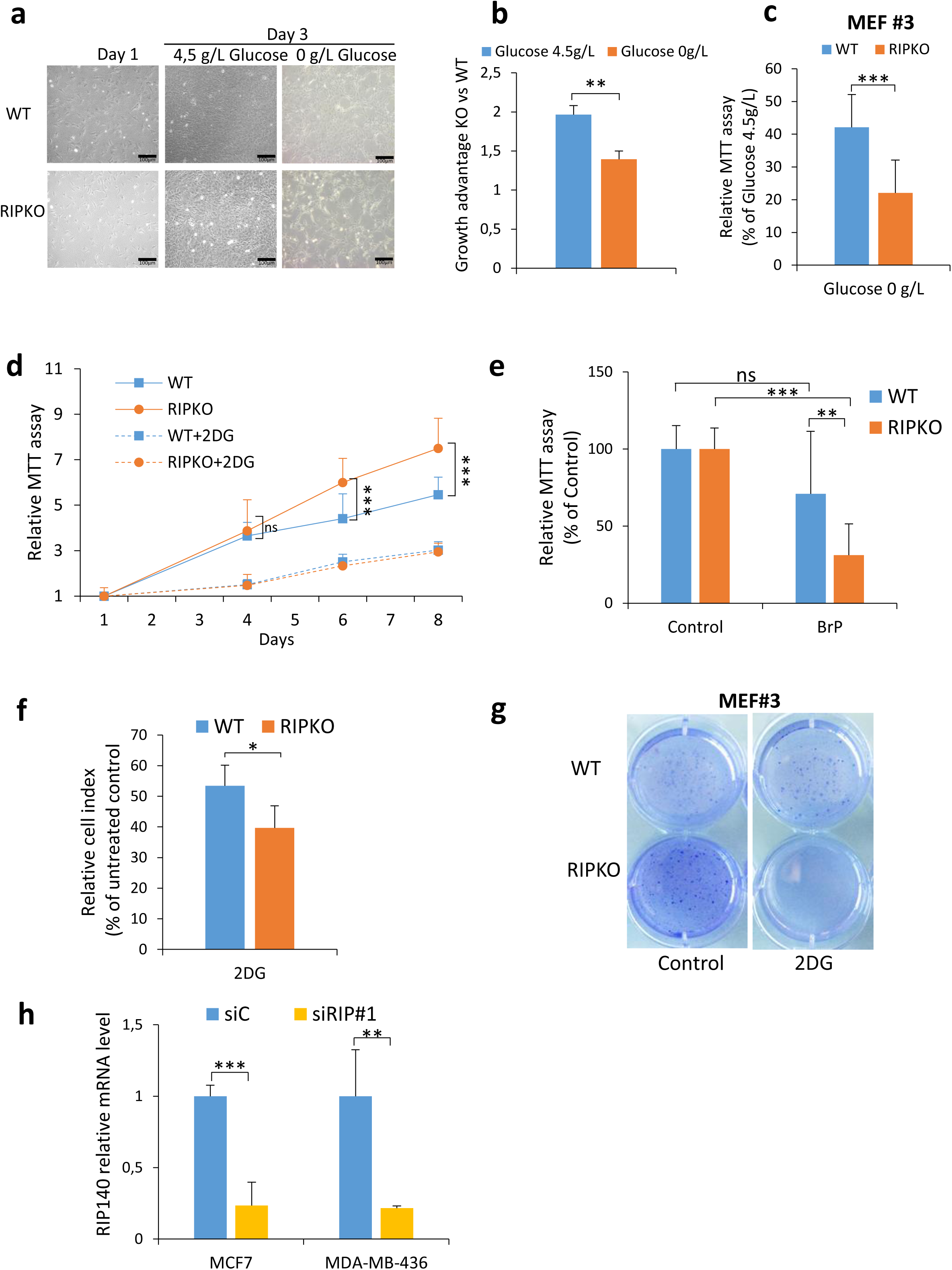
Inhibiting glycolysis reduces the growth advantage of RIP140-deficient cells. Unless otherwise stated, MEF #1 were used. **a**, Representative pictures of MEF #1 under glucose starvation. Scale bars: 100µm. **b**, 3-(4,5-dimethylthiazol-2-yl)-2,5-diphenyltetrazolium Bromide (MTT) assay in MEF #1 cells cultured in DMEM-containing 0g/L or 4,5g/L glucose. The data are normalized to day 1 values and to WT samples representing the growth advantage of RIPKO MEFs over WT MEFs (mean ± SD, n=3, **p <0.01). **c**, 3-(4,5-dimethylthiazol-2-yl)-2,5-diphenyltetrazolium Bromide (MTT) assay in SV40/H-RasV12-transformed MEF #3 cultured in DMEM containing 0g/L or 4.5g/L of glucose for 4 days. Data are normalized to that of cells grown in Glucose 4.5g/L (mean ± SD, n=3, ***p <0.001). **d**, 3-(4,5-dimethylthiazol-2-yl)-2,5-diphenyltetrazolium Bromide (MTT) assay in SV40/H-RasV12-transformed MEF #3 treated or not with 2-deoxyglucose (5mM). The statistical analysis compared the differences in the values between untreated RIPKO and WT samples (mean ± SD, n=3, ***p <0.01, *ns* not significant). **e**, 3-(4,5-dimethylthiazol-2-yl)-2,5-diphenyltetrazolium Bromide (MTT) assay in MEF #1 treated or not with 3-Bromopyruvate (BrP; 100µM) at day 4. Data are normalized to untreated control for each cell line (mean ± SD, n=3, **p <0.01, ***p <0.01, *ns* not significant) **f**, Relative cell index is normalized to untreated control samples for each cell line after four days of 2DG treatment (5mM) (from Fig. 2c; mean ± SD, n=3, *p <0.05, *ns* not significant). **g**, Crystal violet staining of MEF #3 after 3 weeks of 2-DG treatment (5mM) used in Fig. 2f. **h**, RIP140 mRNA expression relative to 28S in MCF7 and MDA-MB-436 cells transfected for 48h with control siRNA (siC) or RIP140 siRNA (siRIP#1) used in Fig. 2h (mean ± SD, n=3, **p <0.01).

**Supplementary Fig. 3.**
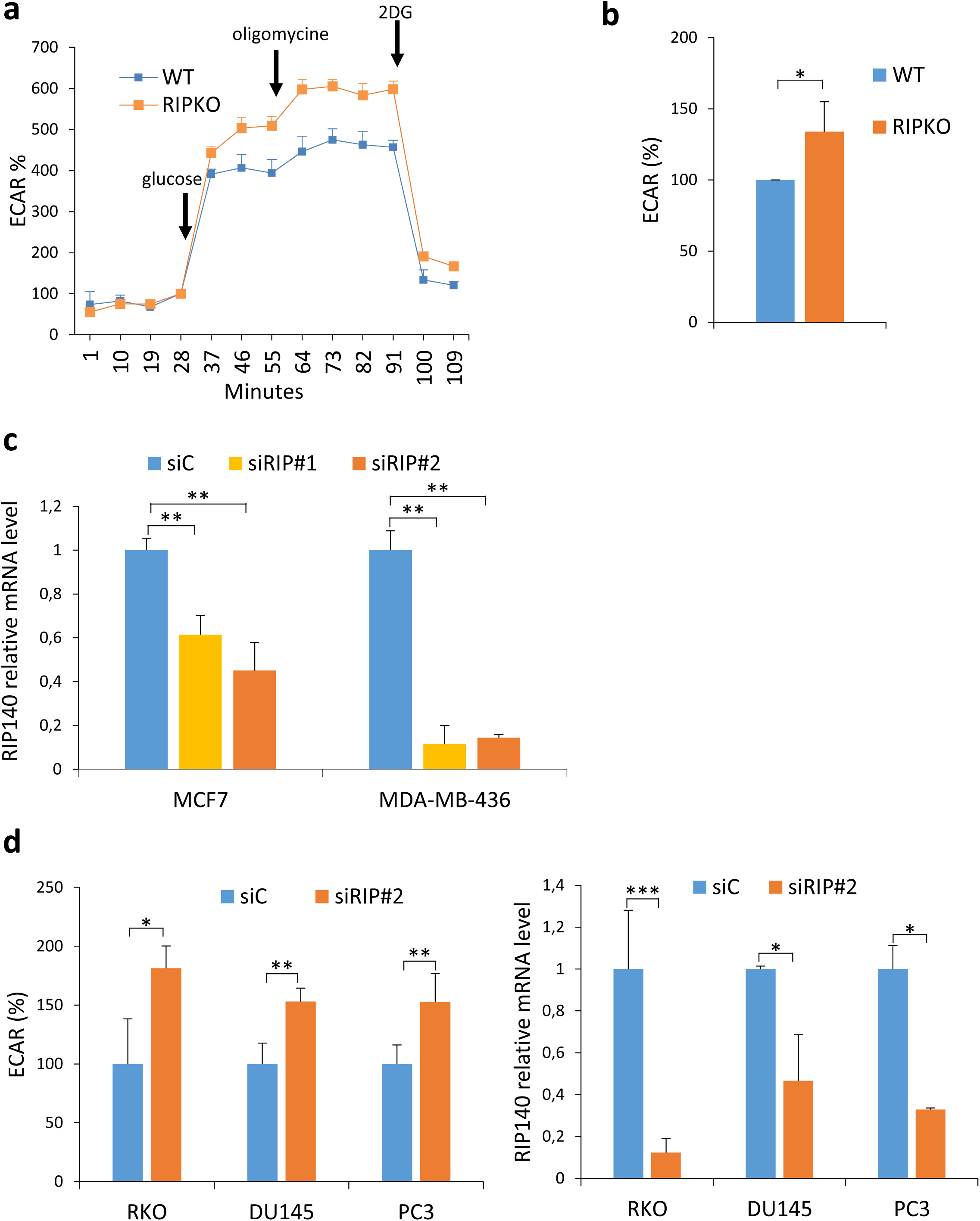
RIP140-deficiency enhances glycolysis. **a**, Measurement of ECAR over time in a representative experiment in MEF #1. Arrows indicate the time points of injection of the different compounds (glucose, oligomycin, 2-deoxyglucose). The values represent the mean of five or six technical replicates and the error bars the standard deviation of the replicate values. Data are normalized to basal measurements and to protein quantity. **b**, ECAR (Extracellular acidification rate) was measured in transformed MEF #3 using the Seahorse XF96 analyzer. Values are normalized to basal measurements and to WT samples (mean ± SD, n=3, *p < 0.05, Unpaired two-tailed t-test). **c**, RIP140 mRNA expression relative to 28S in MCF7 and MDA-MB-436 cells transfected with control siRNA (siC) or two different RIP140 siRNA (siRIP#1, siRIP#2). Values are normalized to control siRNA samples (mean ± SD, n=3, **p < 0.01). **d**, *Left panel*: ECAR (Extracellular acidification rate) was measured in RKO, DU145 and PC3 cells transfected with control siRNA (siC) or RIP140 siRNA (siRIP#2) using the Seahorse XF96 analyzer. Values are normalized to basal measurements and to control siRNA samples (mean ± SD, *p < 0.05, **p < 0.01). *Right panel*: RIP140 mRNA expression relative to 28S in cells used in the left panel. Values are normalized to control siRNA samples (mean ± SD, n=3, *p < 0.05, **p < 0.01).

**Supplementary Fig. 4.**
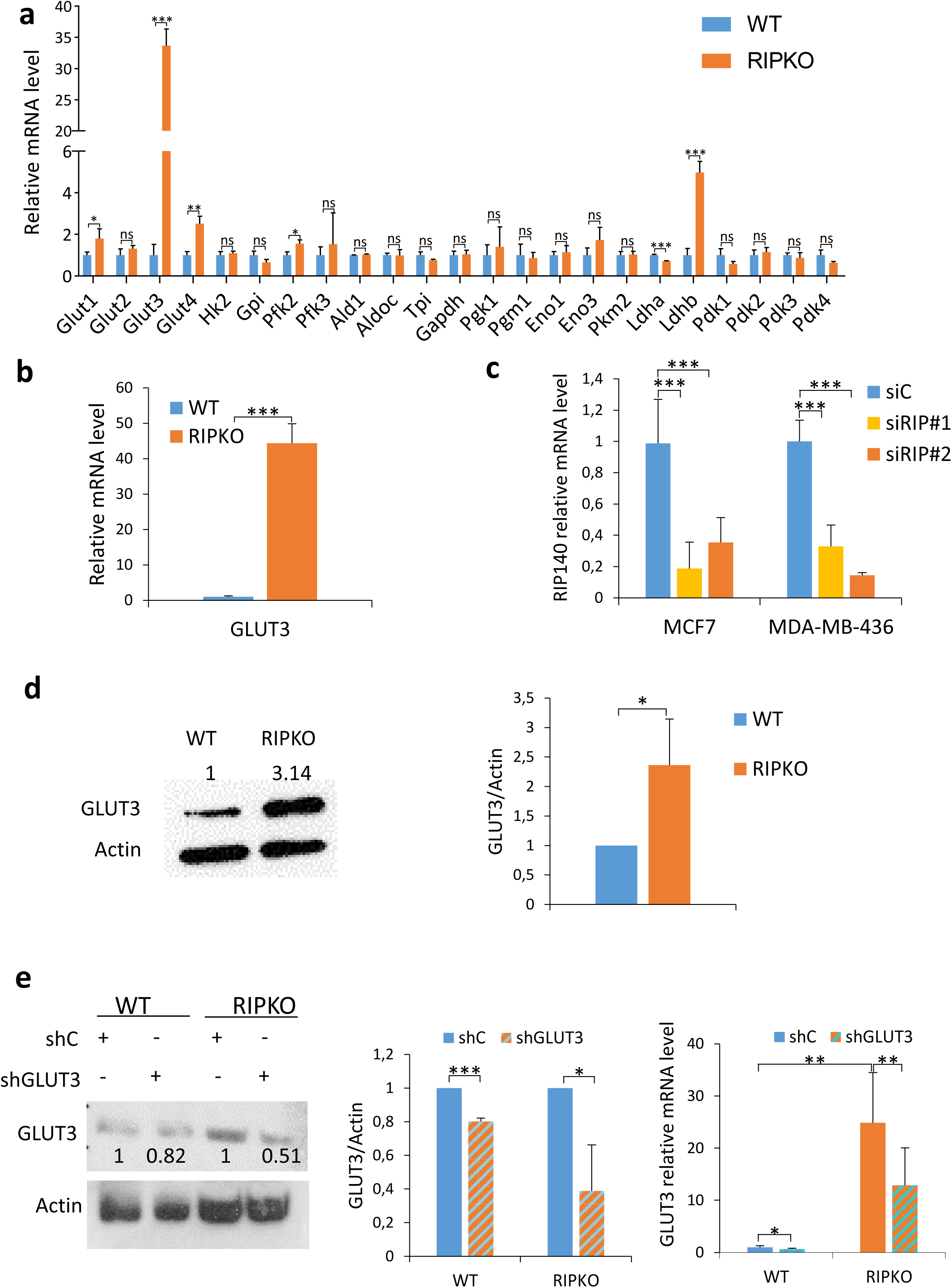
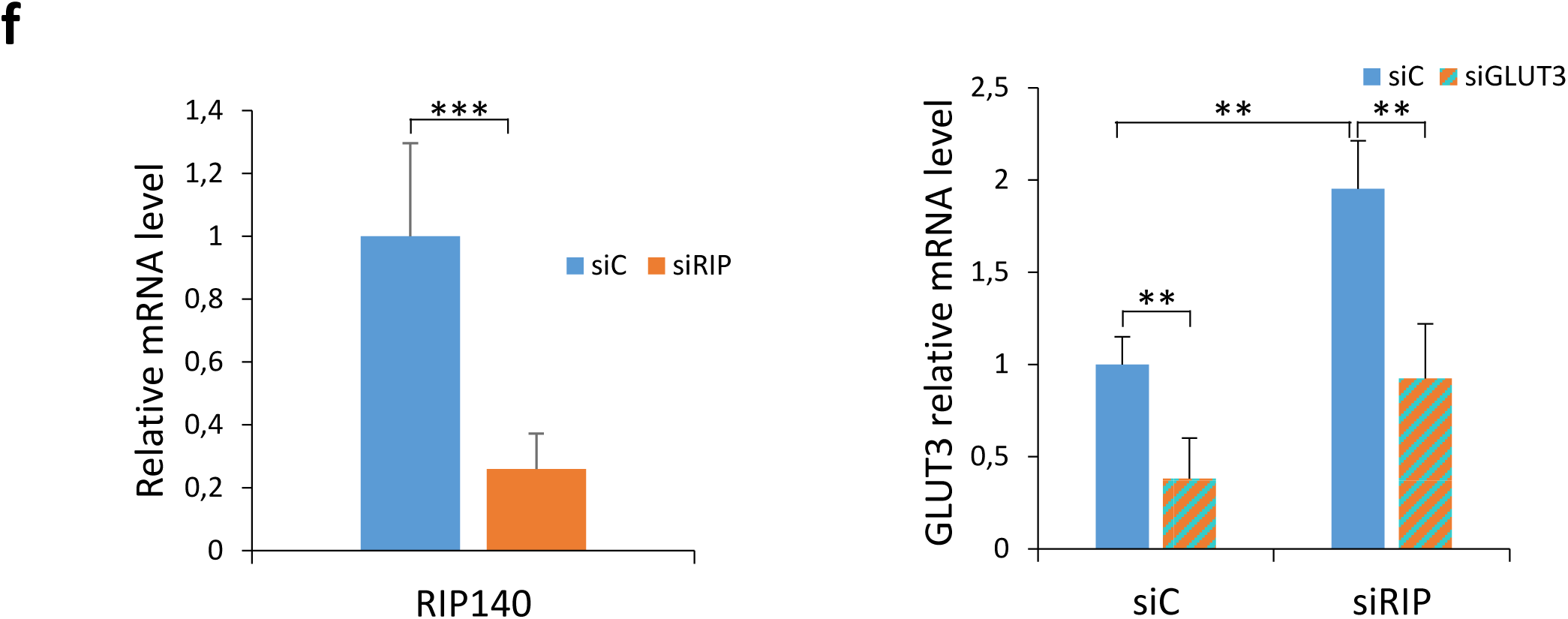
GLUT3 is essential for the growth advantage of RIP140-deficient cells. **a**, mRNA expression relative to RS9 of the indicated genes in MEF #1. Values are normalized to WT samples (mean ± SD, n=3, *p < 0.05, **p < 0.01, ***p < 0.001, *ns* not significant). **b**, Glut3 mRNA expression relative to RS9 in MEF #3. Values are normalized to WT samples (mean ± SD, n=3, ***p < 0.001). **c**, RIP140 mRNA expression relative to 28S in MCF7 and MDA-MB-436 cells after RIP140 downregulation with two different siRNAs (siRIP#1, siRIP#2). Values are normalized to control siRNA (siC) samples (mean ± SD, n=3, ***p < 0.001). **d**, The levels of GLUT3 and Actin were assessed by western blot in MEF #1. A representative blot (left panel) and analysis of band density from three independent experiments (right panel) are shown. Values are normalized to WT samples (mean ± SD, n=3, *p < 0.05). Unprocessed original scans of blots are shown in Supplementary Figure 7. **e**, Knockdown efficiency of Glut3 by specific shRNA was determined by western blot (left panel) and quantified by densitometry (middle panel) and by RT-qPCR (right panel). Values are normalized to WT control shRNA (shC) samples (mean ± SD, n=3, *p < 0.05, **p < 0.01, ***p < 0.001). **f**, RIP140 (left panel) and GLUT3 (right panel) mRNA expression relative to 28S in MDA-MD-436 cells transfected for 48h with RIP140 siRNA (siRIP) combined with GLUT3 siRNA. Values are normalized to control siRNA (siC) samples (mean ± SD, n=3, *p < 0.05, **p < 0.01, ***p < 0.001).

**Supplementary Fig. 5.**
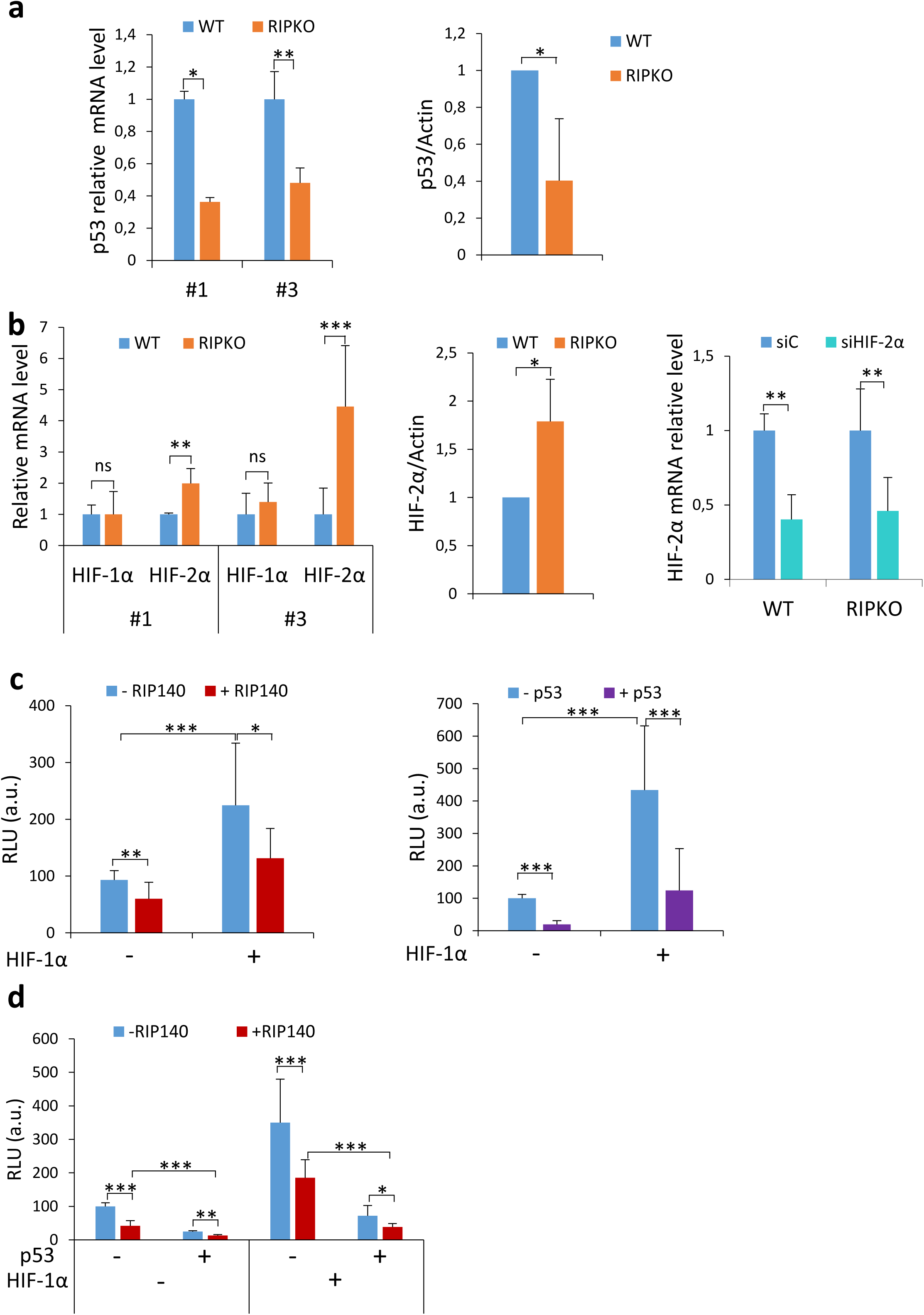
RIP140 and p53 interact to inhibit the expression of GLUT3. **a**, *Left panel:* p53 mRNA expression relative to RS9 in MEF #1 and #3 cells. Values are normalized to WT samples (mean ± SD, n=5, *p < 0.05, **p < 0.01). *Right panel:* Densitometry analysis of p53 protein expression assessed by western blot and normalized to the housekeeper Actin (mean ± SD, n = 3, * p <0.05). **b**, *Left panel*: HIF-1α and HIF-2α mRNA expression relative to RS9 in MEF #1 and #3 cells. Values are normalized to WT samples (mean ± SD, n=5, **p < 0.01, ***p <0.001, *ns* not significant). *Middle panel*: Densitometry analysis of HIF-2α protein expression assessed by western blot and normalized to the housekeeper Actin (mean ± SD, n = 3, * p <0.05). *Right panel*: HIF-2α mRNA expression relative to RS9 in MEF #1 cells transfected with control siRNA (siC) or HIF-2α siRNA (siHIF-2α). Values are normalized to control siRNA samples (mean ± SD, n=3, **p < 0.01). **c**, Luciferase activity assay in MEF #1 RIPKO cells transfected with pcDNA3.1-HIF-1α plasmid, the luciferase reporter containing hypoxia inducible response elements and the luciferase reporter TK-Renilla and pef-cMyc-RIP140 (left panel) or pcDNA3.1-p53 (right panel). Luciferase values were normalized to the Renilla luciferase control and to the values of samples transfected with empty vectors (mean ± SD, n=3, *p <0.05, **p <0.01, ***p <0.001). **d**, Luciferase activity assay in MEF #1 RIPKO cells transfected with pcDNA3.1-HIF-1α plasmid, the luciferase reporter containing hypoxia inducible response elements and the luciferase reporter TK-Renilla, pef-cMyc-RIP140 and pcDNA3.1-p53 vectors. Luciferase values were normalized to the Renilla luciferase control and to the values of samples transfected with empty vectors (mean ± SD, n=3, *p <0.05, **p <0.01, ***p <0.001).

**Supplementary Fig. 6.**
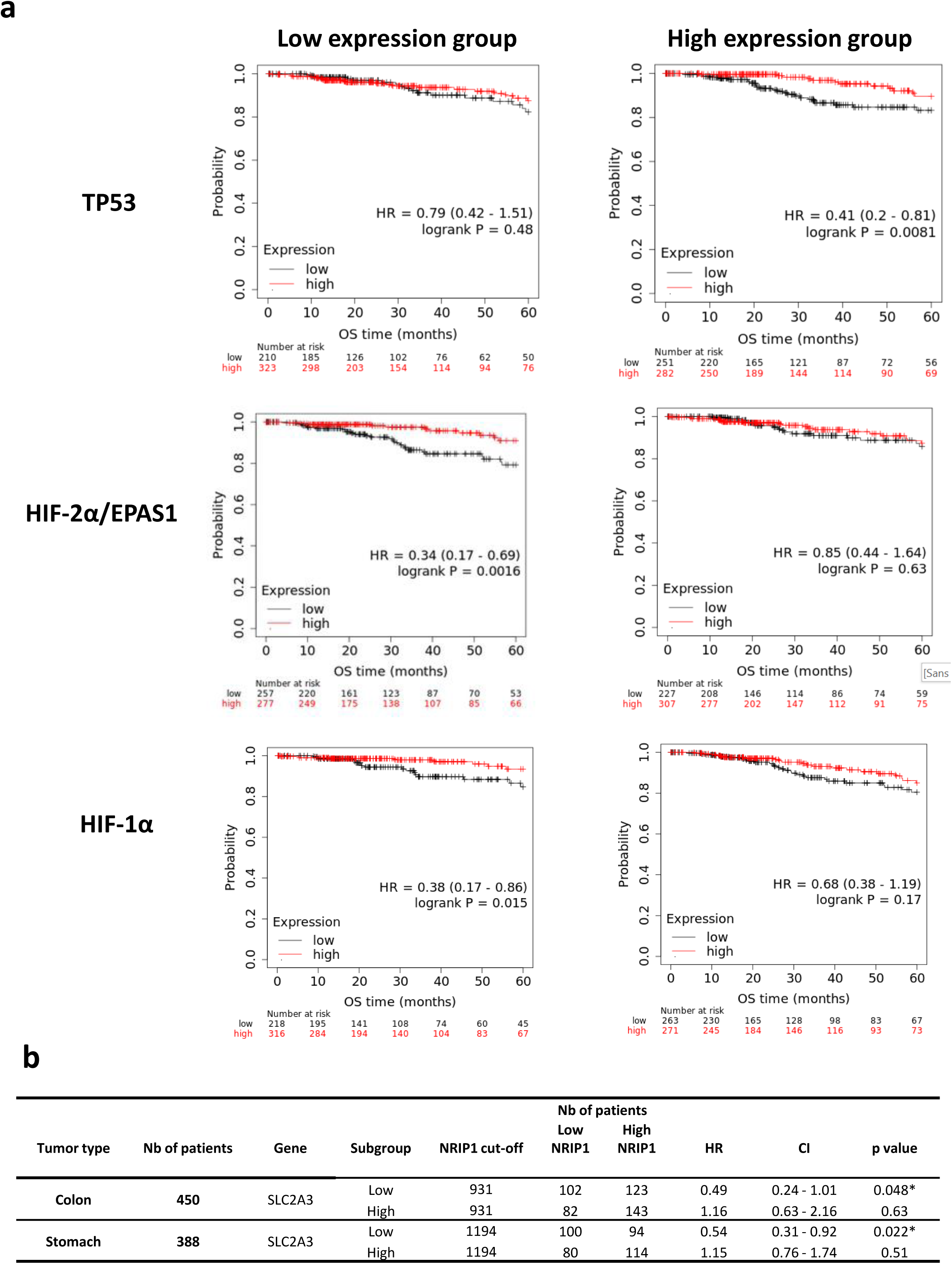
The prognostic value of RIP140 is correlated with the levels of GLUT3 expression. Kaplan-Meier of 1068 cases of breast cancer with statistical significance being assessed using the log-rank test from the TCGA dataset. **a**, Groups have been defined on the basis of the median p53 (TP53, top panel), HIF-2α (EPAS1, middle panel) and HIF-1α (HIF1A, bottom panel) expression. Kaplan-Meier analyses plotting the survival curved used to define RIP140 prognostic value in Fig. 6c. **b**, GLUT3 (SLC2A3) expression groups have been defined on the basis of the median GLUT3 expression for colon and stomach cancer patients. RIP140 prognostic value was significantly associated with better overall survival in low GLUT3 expression group (colon, P=0.048; stomach, P=0.022).

**Supplementary Fig. 7.**
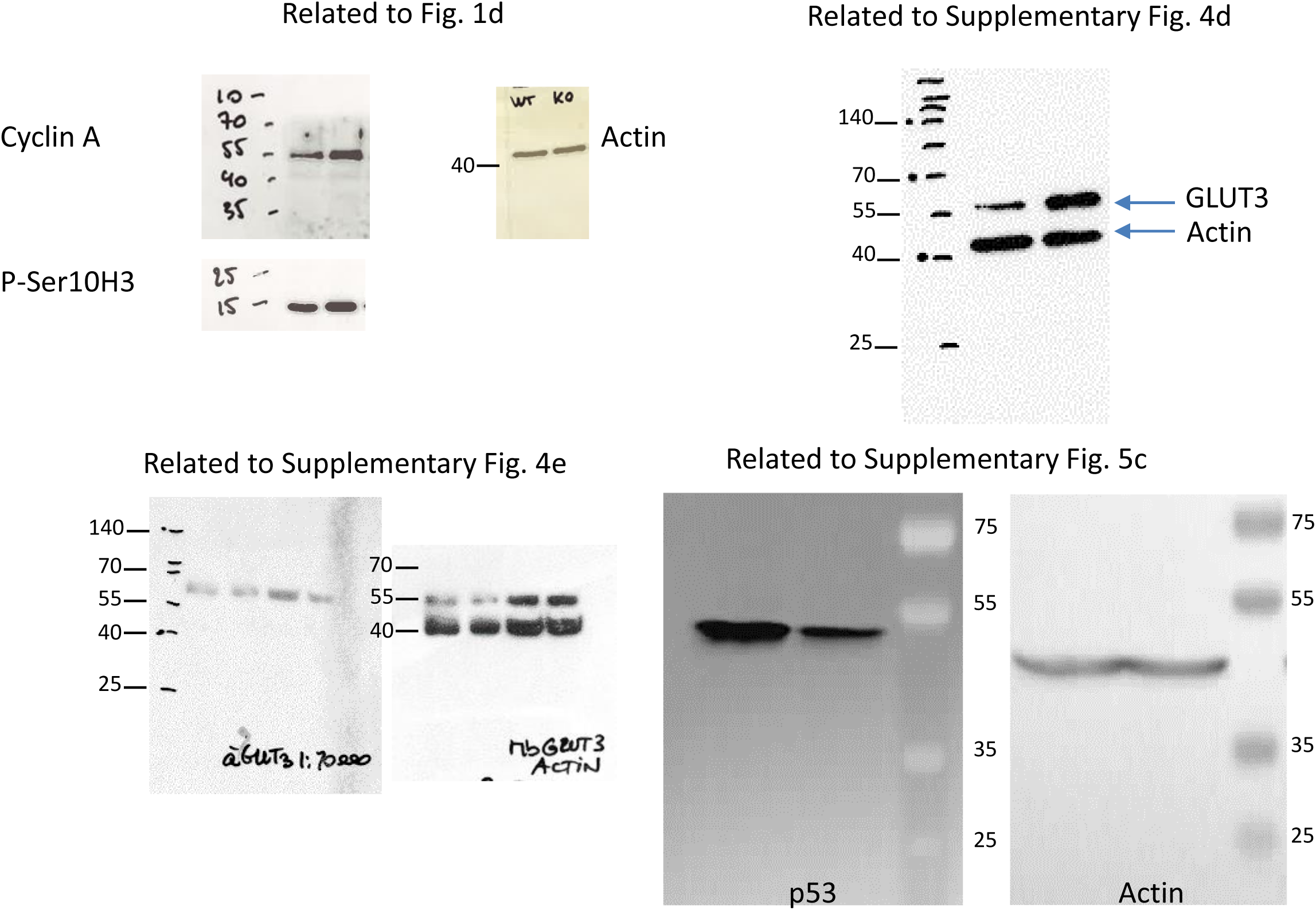
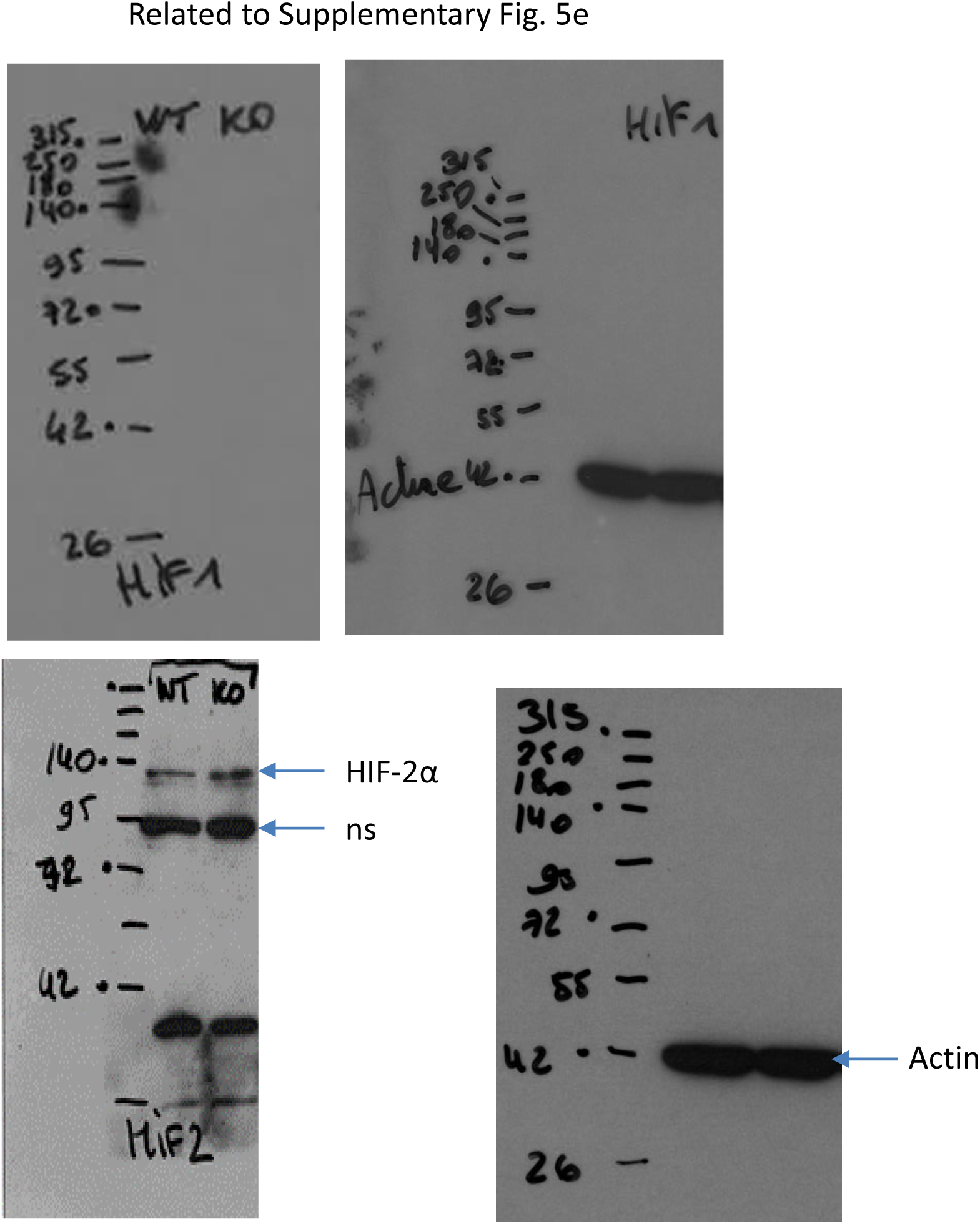
Unprocessed original scans of blots. Unprocessed images of all Western blots as indicated. Molecular size markers in kDa.

